# Deep learning identifies and quantifies recombination hotspot determinants

**DOI:** 10.1101/2021.07.29.454133

**Authors:** Yu Li, Siyuan Chen, Trisevgeni Rapakoulia, Hiroyuki Kuwahara, Kevin Y. Yip, Xin Gao

**Affiliations:** Department of Computer Science and Engineering, The Chinese University of Hong Kong, Hong Kong SAR, China; Computational Bioscience Research Center, Computer, Electrical and Mathematical Sciences and Engineering Division, King Abdullah University of Science and Technology (KAUST), Thuwal 23955-6900, Saudi Arabia; The CUHK Shenzhen Research Institute, Hi-Tech Park, Nanshan, Shenzhen 518057, China; Max Planck Institute for Molecular Genetics, Ihnestrasse 63-73, 14195 Berlin, Germany

## Abstract

Recombination is one of the essential genetic processes for sexually reproducing organisms, which can happen more frequently in some regions, called recombination hotspots. Although several factors, such as PRDM9 binding motifs, are known to be related to the hotspots, their contributions to the recombination hotspots have not been quantified, and other determinants are yet to be elucidated. Here, we develop a computational method, RHSNet, based on deep learning and signal processing, to identify and quantify the hotspot determinants in a purely data-driven manner, utilizing datasets from various studies, populations, sexes, and species. In addition to being able to identify hotspot regions and the well-known determinants accurately, RHSNet is sensitive to the difference between different PRDM9 alleles and different sexes, and can generalize to PRDM9-lacking species. The cross-sex, cross-population, and cross-species studies suggest that the proposed method has the potential to identify and quantify the evolutionary determinant motifs.

**Teaser:** RHSNet can accurately identify and quantify recombination hotspot determinants across different studies, sexes, populations, and species.

## Introduction

The Recombination is an essential and fundamental genetic process in meiosis, which introduces new combinations of alleles and generates haplotypic diversity in sexually reproducing organisms, driving evolution and biodiversity(*1–3*). Although the molecular mechanism of this process has not been fully uncovered(*1*), it is believed that in many species, including humans and mice, the event begins with the binding of DNA by the histone methyltransferase PRDM9(*4*). The double-strand break (DSB) machinery, including the meiotic topoisomerase-like protein SPO11(*5*), is then recruited by an unknown mechanism, forming DSBs. Specialized pathways repair these breaks, with the majority leading to noncrossovers (NCOs) while the minority developing crossovers(*6*) (COs). Although PRDM9 binds ubiquitously throughout the genome, the distribution of COs is nonrandom, clustered in narrow regions, called recombination hotspots(*1, 4*). Despite the unclear reason for forming hotspots, the following factors are suggested to be related to the locations of hotspots(*2, 7*). The DNA binding domain of PRDM9 influences sequence specificity and the formation of DSBs(*4, 8–10*); histone modifications can influence the chromatinic local structure and thus affect crossover formation(*3, 11*); recombination occurs more frequently in GC-rich regions(*2, 12*). Yet, more factors influencing the recombination events and hotspot formation, and the molecular mechanism behind it are yet to be discovered(*1, 13*).

Several studies(*2, 6-8, 13-18*) have been conducted to demystify this essential genetic process, resulting in a large amount of data with different properties. The availability of such datasets enlightens the possibility of investigating this problem from a different angle, that is, in a data-driven manner. Despite the extensive biological experiments and the development of computational tools to construct genetic maps(*3, 19*) and perform binary classification(*20–22*) of hotspot and coldspot sequences, computationally, researchers have not investigated the determinants of recombination hotspots systematically and quantitatively. Based on the accumulated datasets from the previous studies, it is very promising to develop computational methods to perform cross-study, cross-sex, cross-population, and cross-species investigation, potentially providing more insights into this crucial biological process.

Deep learning has been proven to be a successful approach for performing classifications(*23, 24*). However, directly applying deep learning models to this problem may cause difficulties in studying the recombination hotspot determinants quantitatively due to the complexities and interpretability issue of the model(*25*). To analyze the accumulated recombination data and facilitate the study of the yet unclear recombination process, we propose a novel transparent computational method, RHSNet, which combines the strength of deep learning(*24*), activation backpropagation(*26*), and signal processing(*27*), to systematically identify and quantify the recombination hotspot determinants taking advantage of data from multiple previous studies crossing different populations(*2, 3, 11, 13*), sexes(*17*), and species(*6, 11*). In addition to predicting hotspot sequences accurately and identifying the well-known determinants, our method can quantify the relative contribution of each determinant, showing their differences in different sexes and species, as well as their evolution across different populations.

## Results

### Accurate, flexible, and interpretable identification of recombination hotspots

Our method leverages the strength of deep learning, activation backpropagation, and signal processing to first predict the recombination hotspot sequences, then quantify the contribution of the input information, and finally extract determinants, such as the PRDM9 binding motif. As shown in Fig. 1, during the prediction, the input sequences of various lengths, depending on the data (Materials and Methods), go through a specific deep learning model, which consists of two 1-D convolutional layers as the sequence feature extractor, a Gated Recurrent Unit (GRU) for capturing long-range information, and a multi-head attention layer for detecting interactions within the sequence (Fig. S2), to output useful information from the raw sequences. Because histone modifications are also shown to be crucial to recombination(*3, 11*), we use another deep learning module to process the information, including H3K4me3 and H3K36me3 from testis and ovary. Then, all the above information is combined to predict whether the input sequence is a hotspot sequence. In addition, we are further interested in identifying and quantifying the recombination hotspot determinants. One previous study(*22*) extracts motifs by considering only the activation of the first layer in the deep learning model. But this method omits the complexity of the downstream layers and has difficulty in quantifying the motif’s contribution based on the entire model. To resolve the issue, we utilize an activation backpropagation method(*26*). The prediction of a specific sequence is backpropagated through the entire network back to the original inputs to assign contribution scores to the motifs. Note that we consider the entire deep learning model and compute the score in a purely data-driven manner. However, extracting determinant information remains a problem because the computed scores can be noisy, with peaks having various lengths along the sequence, as shown in Fig. 1. We resolve this problem using signal processing techniques. We apply a low-pass filter onto the contribution score array. Then, we extract the significant motifs between two valleys with a peak. The user has the freedom to choose the low-pass filter, either obtaining long determinants or short ones with high confidence. Based on the outputs of RHSNet, we further perform comprehensive quantitative analysis, as shown in Fig. 1, which will be discussed in detail below.

**Fig. 1.**
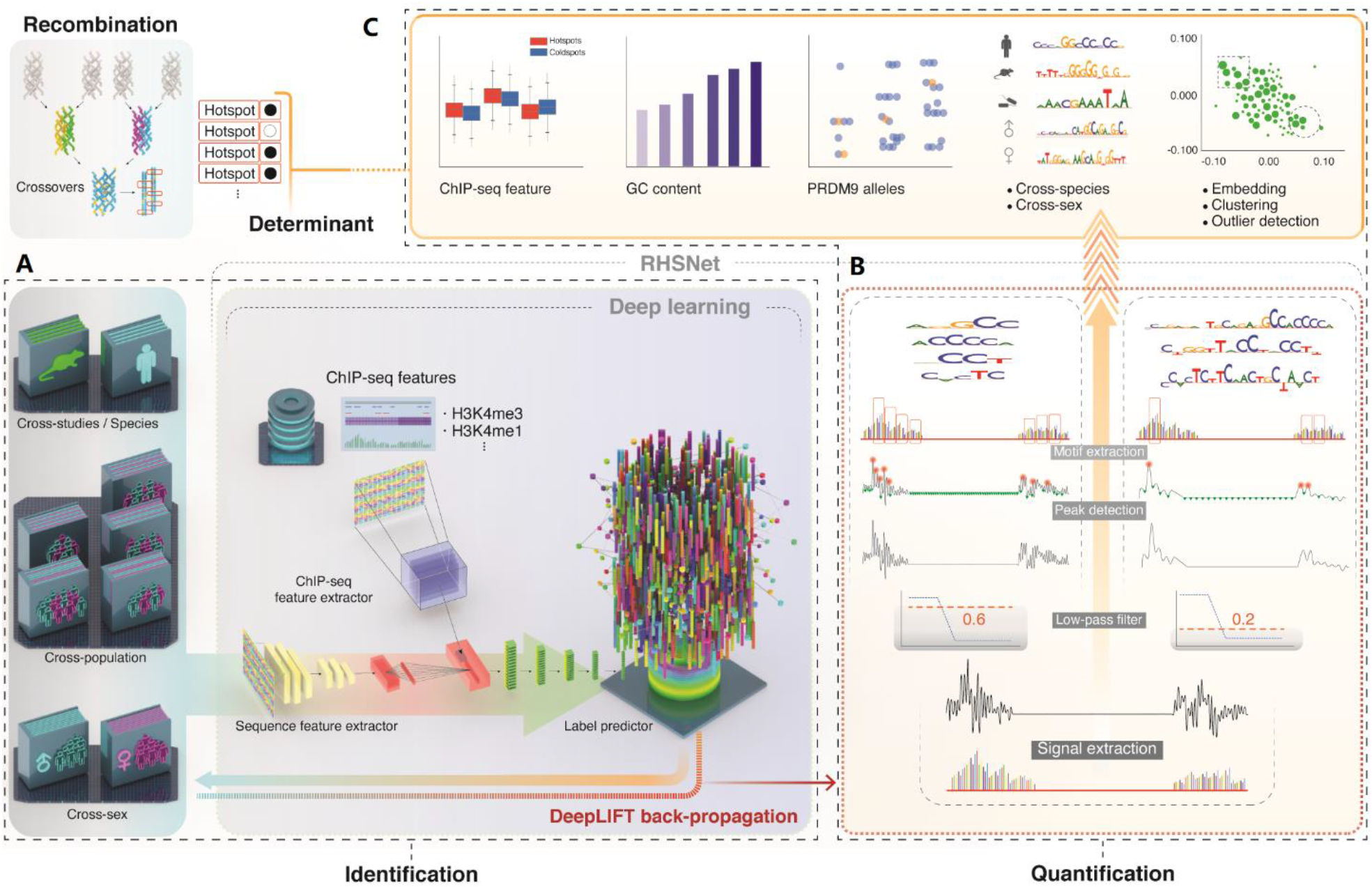
Overview of the proposed framework, RHSNet, along with the proposed filter-based motif extraction approach. **(A)**: The deep learning algorithm of RHSNet accurately identifies recombination hotspots from different studies/species/populations/sexes, considering the ChIP-seq information. The model consists of two 1-D convolutional layers as the sequence feature extractor, a Gated Recurrent Unit (GRU) for capturing long-range information, and a multi-head attention layer for detecting interactions within the sequence. **(B)**: RHSNet has a low-pass filter-based motif extractor that can quantify the contribution of hotspot-determinant motifs with flexible lengths (4bp-30bp). We propose such a method based on gradient back-propagation and signal processing. **(C)**: Comprehensive analysis across different studies/species/populations/sexes, based on RHSNet, provides more insights into the biological process and suggests the effectiveness of the proposed method.

Although our method is not designed specifically to perform binary recombination hotspot prediction, it can achieve superior performance on the task, compared with the previous methods. Here, we report the performance of the classification module in RHSNet (the Identification module in Fig. 1). We evaluate the proposed deep learning model’s performance on different datasets (HapMap II(*28*), Icelandic(*2*), and Sperm(*13*)) (Materials and Methods), different sexes(*2*), different populations(*28*), different species(*6, 11*) and across different evaluation criteria, whose results are shown in Fig. 2. With the same input data and evaluation criteria, our deep learning model in RHSNet is constantly better than the competing models, including convolutional neural network, Equivariant CNN(*22*) and PseDNC(*20*) (Table S5), across different conditions in terms of F1 score (Fig. 2A), except for the paternal Icelandic dataset. Meanwhile, on the sex-specific dataset, for which we can extract ChIP-seq information from the corresponding ovary and testis tissues, we utilize six histone modifications (H3K4me1, H3K4me3, H3K27ac, H3K9me3, H3K36me3, H3K27me3) from testis and ovary tissues for the paternal and maternal datasets, respectively. Adding such information into our model (RHSNet-chip) can further boost the deep learning model’s performance, which is consistent with the previous research(*3, 11*). To further test the robustness of our model, we evaluate the model’s performance on hotspot regions with various lengths (Fig. 2B, on the Icelandic dataset), using a different evaluation criterion, Matthews Correlation Coefficients (MCC). As illustrated in Fig. 2B, our method is consistently better and shows more stable and robust performance than the baseline methods across different resolutions. As the resolution goes down and the prediction becomes less demanding, all the models’ performance improves, although the training dataset size decreases. To further validate our method, we test it against a dataset from a different species, the Mice dataset(*11*). Evaluated on the dataset with Area Under the Receiver Operating Characteristic curve (AUROC), RHSNet can significantly improve over the existing methods (Fig. 2C). More statistical results about the identification performance can be referred to (Supplementary Materials).

**Fig. 2.**
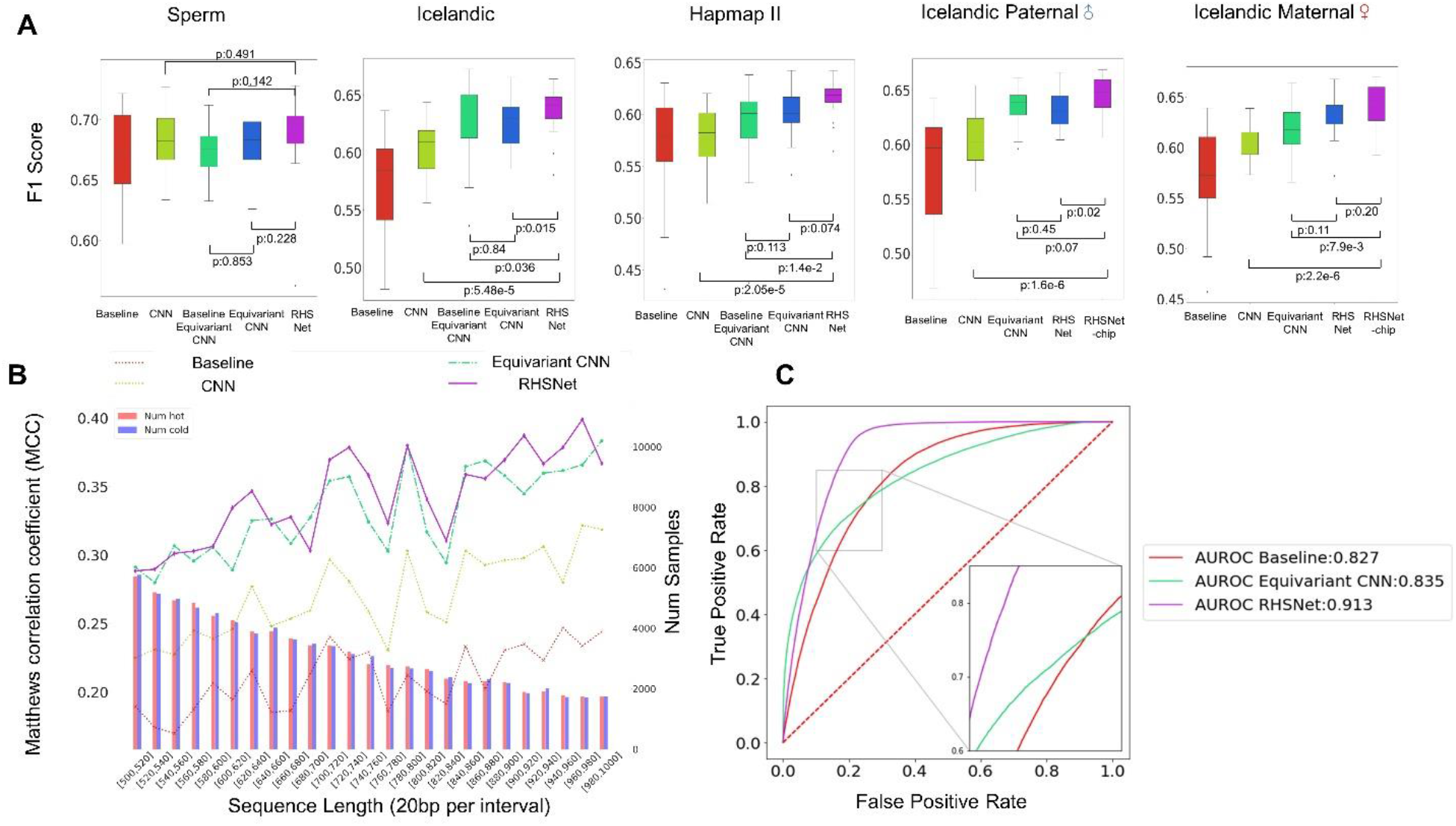
Performance of RHSNet across different studies/sexes/species. Notice that, here, RHSNet refers to the Identification module of RHSNet in Fig. 1. **(A)** The performance of RHSNet on datasets from different studies and different populations. We used 5-fold cross-validation to evaluate the performance of different methods. The box plots show the F1-Score distributions from four trials of the 5-fold cross-validation. P-values calculated from the two-tailed Student’s t-test indicate the significance of the improvement. RHSNet-chip refers to RHSNet accompanied with ChIP-seq information. We used several histone modifications from testis for the male dataset and ovary for the female dataset. **(B)** Robustness testing with Matthews correlation coefficients (MCC) on the Human Icelandic dataset of RHSNet over different input lengths ranging from 500bp to 1000bp. The number of hotspot and coldspot sequences within each interval of 25bp is showed as a histogram at the bottom. RHSNet’s performance is robust across different resolutions. **(C)** On the Mice dataset, RHSNet also shows significant performance improvement on predicting the hotspot sequences in terms of the AUROC score. This result suggests the generalization ability of our method on different datasets, even across different species.

### PRDM9 binding motif, GC content, and histone modification affect recombination hotspots

The existing research has shown that PRDM9 binding motif(*4, 8–10*), histone modification(*3, 11*), and GC content(*2, 12*) influence the recombination hotspot. We use RHSNet to analyze how the above factors are related to the recombination hotspots in different datasets. As we have discussed, in our method, with different low-pass filter factors, we can extract motifs with different lengths and different enrichment factors (Fig. 1, Fig. 3A, Materials and Methods). In Fig. 3B, we compare the top-ranked motifs regarding the enrichment factor from different datasets (HapMap II(*28*), Icelandic(*2*), and Sperm(*13*)) with different filter factors against the PRDM9 binding motif^4^: *CCNCCNTNNCCNC* and SPO11-oligo(*11*). Clearly, in the Icelandic dataset, the top-ranked motifs are highly correlated (top10: 87.5% ± 1.01; top100: 64.1% ± 1.54) with the canonical PRDM9 motif regarding the pairwise sequence alignment matching score. Although, compared with Icelandic, the PRDM9 pattern is less significantly enriched in the top-ranked motifs from the HapMap II dataset, we still obtain 53.91% GC content in the top 100 motifs (Fig. 3D), with the PRDM9 binding pattern appearing in these motifs. In contrast, the top-ranked motifs from the Sperm datasets are different from the ones from the other two datasets, though the PRDM9 pattern still appears. Unlike the HapMap II(*28*) and Icelandic(*2*) datasets, the Sperm(*13*) dataset focuses more on comparison across individuals’ cells rather than aggregating them, and is resolved to much larger regions, with the median resolution as 240 kilo-base pairs (kbp), among which 9,746 (1.2%) are inferred within 10 kbp. Consequently, we inevitably involve noisy sequences in the training dataset, which reduces the sensitivity of our method and also leads to lower prediction confidence compared to the other datasets when the filter factor is 0.4.

**Fig. 3.**
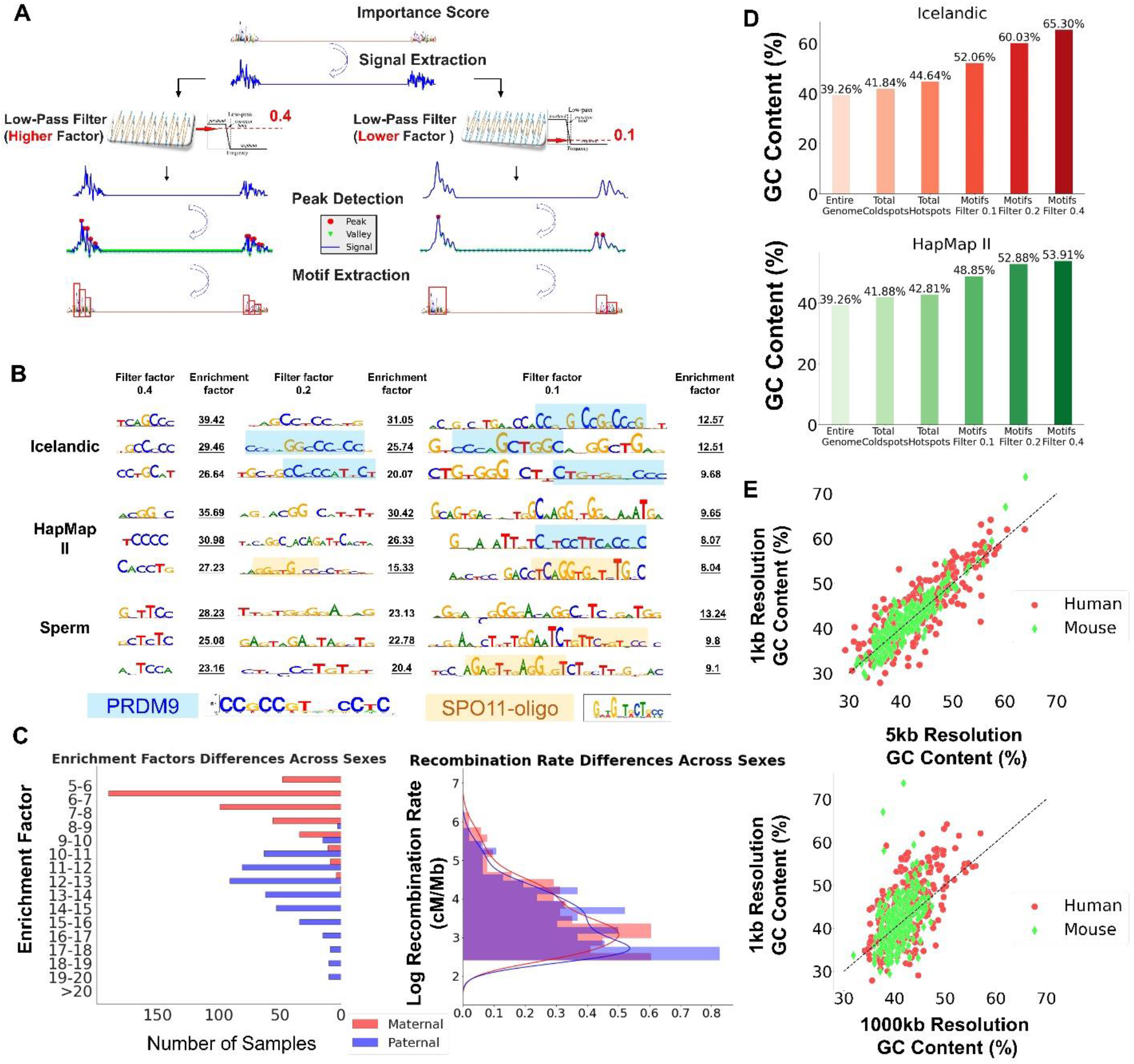
RHSNet quantifies the contribution of PRDM9 binding motifs in variant lengths across different studies/populations/sexes. **(A)** RHSNet can extract motifs of variant lengths with different low-pass filter factors. **(B)** Assembled motif detection results are shown using different low-pass filters ranging from 0.1 to 0.4. PRDM9 binding motifs and 12-bp motif enriched in SPO11-oligo hotspots show high enrichment factors. **(C)** The logged recombination rates and the enrichment factor distribution across sexes within the detected motifs. The recombination rates are higher in the females, while the enrichment factors are higher in the males. The lower erosion rate of the hotspot motifs in males may make the event determinants more conservative. **(D)** GC content compared across different studies. We show the GC content among the entire genome, total hotspots, total cold spots, and RHSNet’s detected motifs. **(E)** GC content of recombination hotspots cropped in different resolutions (1kbp, 5kbp, 1000kbp) across different species. The GC content is usually higher in the hotspots and the determinant motifs compared to the nearby regions.

GC content is shown to be positively correlated with the recombination rate(*2, 12*). As shown in Fig. 3D and Fig. S6, in all the datasets, the GC content of the hotspots is indeed higher than that of the entire genome (HapMap II: 44.64% VS 39.26%, Icelandic: 42.81% VS 39.26%, Sperm: 48.16% VS 39.26%). The comparison of GC content in hotspots across different resolutions (Fig. 3E) also suggests that the determinants in the central hotspot area are GC-richer than the marginal. Interestingly, the GC content of the coldspots (40.72%∼41.88%), where the recombination rates are the lowest in the genome, is not lower than that of the entire genome, although it is lower than that of the hotspots. For some datasets, such as HapMap II, the result is expected because the coldspot set was constructed to match the GC content of the hotspot one. However, for the other datasets, the coldest coldspots show similar GC content, which suggests that GC content itself may not be the causation of hotspots. Instead, the higher GC content in hotspots may be the consequence of the determinant motifs, which are GC-rich, such as the PRDM9 binding motif. In the HapMap II dataset and the Icelandic dataset, where the resolution is high enough (up to 642bp), the determinant motifs identified by our model have much higher GC content than that of the overall hotspot regions. Furthermore, as we increase the filter factor, which forces our method to output shorter motifs with higher enrichment factors, the GC content increases further in these motifs (up to 65.3%). The separated paternal map and maternal map show a similar trend as the average map in the Icelandic dataset (Fig. S7), although the signal is reduced because fewer people are included in each map. Intuitively, the results are consistent with the previous discoveries, as the PRDM9 motif, which is GC-rich, is the most popular motif in the hotspot region.

Histone modifications usually accompany the recombination event, and in the PRDM9 knockout organism, DSB is directed by histone modifications(*29*) (Fig. 4A). We quantify the contribution of different histone modifications to recombination hotspot formation using activation backpropagation. In Fig. 4B, on the Icelandic dataset, we compare the distribution of different features from three modifications between hotspots and coldspots, including H3K4me1, H3K4me3, and H3K36me3. The distribution differences suggest that the histone modification patterns are different in the two kinds of regions. We use the Icelandic maternal data and the histone modifications from the ovary to further investigate such features. In Fig. 4C, we show the feature correlations among six histone modifications across hotspots. H3K4me1is correlated with H3K36me3, which is similar to that in the functional elements in the genome(*30*). To illustrate the contribution of histone modification features, in Fig. 4D, we visualize the 2D vector embedding of recombination hotspots and coldspots from the Principal Component Analysis (PCA) extracted from the last layer of CNN, RHSNet, and RHSNet-chip. As shown in the figure, the proposed deep learning model in RHSNet can learn different representations for hotspots and coldspots, and thus identify the hotspot regions. RHSNet-chip, incorporating the histone modification information, can further enlarge the difference in the learned representation between hotspots and coldspots. To further quantify the contribution of features from histone modifications, we use activation backpropagation across the entire network, visualizing their importance scores in Fig. 4E. The results are consistent with previous studies(*29*), with H3K4me3 and H3K36me3 being the two most essential modifications. Other related modifications, such as H3K4me1(*31*) and H3K27ac(*32*), are also captured by our method, although they are less studied for this problem.

**Fig. 4.**
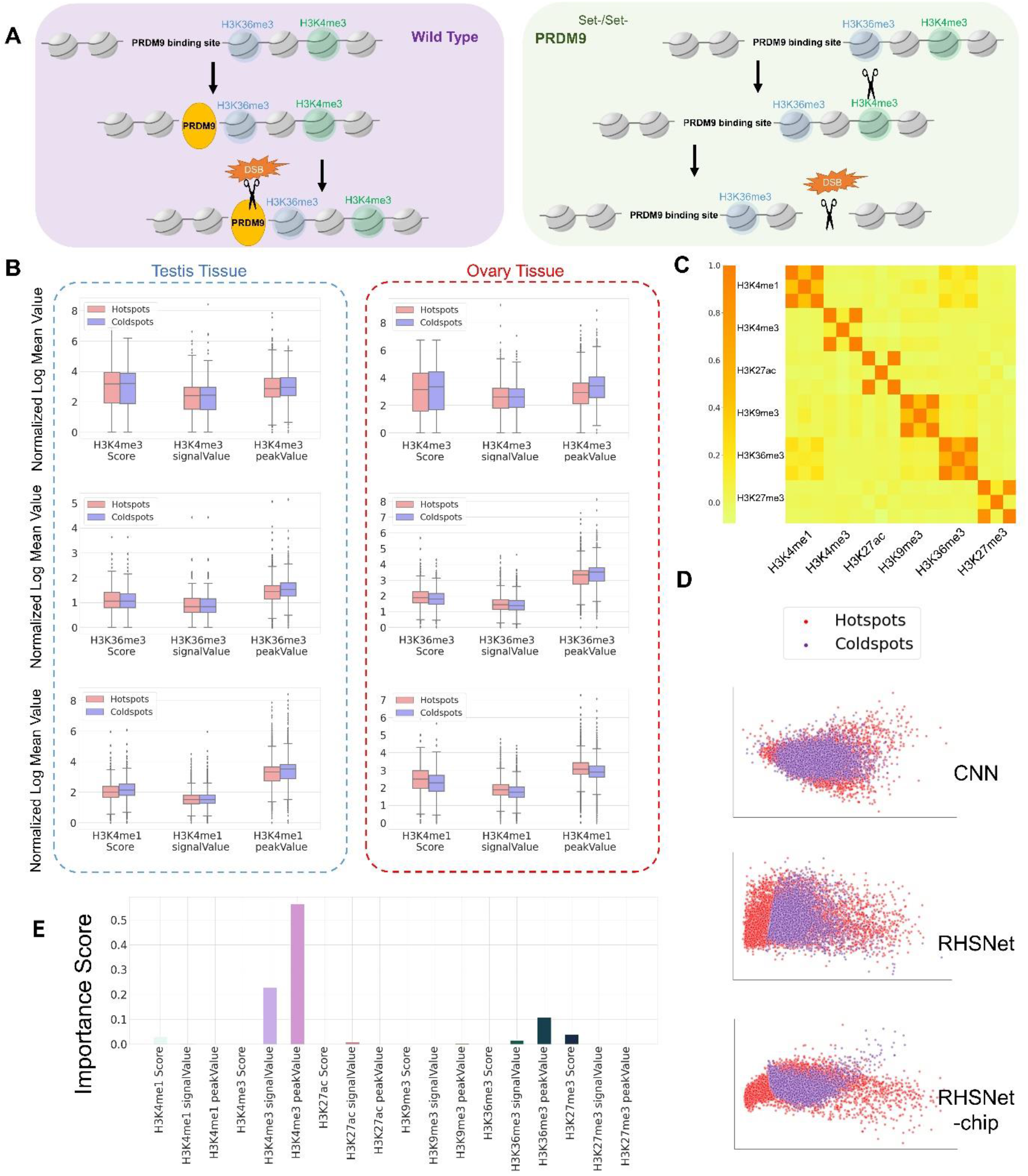
Histone modification affects recombination hotspots formulation. **(A)** The DSB formation machinery (scissors) is directed to PRDM9 binding sites with functional PRDM9 protein. However, in the absence of PRDM9, DSB is directed to PRDM9-independent H3K4me3 marks. **(B)** On the maternal and paternal maps from the Icelandic dataset, we show the feature distribution comparison from recombination hotspots and coldspots over three histone modifications: H3K4me1, H3K4me3, and H3K36me3. **(C)** Heatmap of the eighteen feature correlations within six histone modifications across hotspots. **(D)** For the ChIP-seq feature of female adult’s ovary tissue, we show the 2D vector embedding of recombination hotspots and coldspots from the Principal Component Analysis (PCA) extracted from the last layer of CNN, RHSNet, and RHSNet-chip. The difference between hotspot features and coldspot features is more significant in the RHSNet-chip framework, demonstrating the importance of the ChIP-seq feature, although RHSNet alone is significant enough. **(E)** The importance score calculated from the contribution back-propagation in the ChIP-seq feature extraction branch quantifies the contribution of each histone feature to hotspot formulation. The activation of the H3K4me3 features and the H3K36me3 features suggests a higher contribution to the recombination event.

### RHSNet reveals the contribution of different PRDM9 alleles in different populations

Not only has PRDM9 been found to be the major determinant of the recombination hotspots in humans and mice(*4*), but different PRDM9 alleles are also believed to influence recombination hotspot activities in humans(*3, 18, 33*). PRDM9-A is the most abundant allele in human populations (found in around 86% of European and around 50% of African populations), and PRDM9-C is the second most common one in African populations (12.8%)(*33*). The two alleles have different binding preferences (Materials and Methods). Despite the imperfect way of identifying the motifs, PRDM9-C binding motifs are found to potentially elevate the recombination rates in the African populations. On the dataset from phase 3 of the 1000 Genomes Project(*18*), in the African population, both detected PRDM9- A/C binding motifs (Materials and Methods) show significantly higher recombination rates than the other populations (Fig. 5A), which is consistent with the previous study(*3*). Furthermore, among the top 100 motifs (low-pass filter: 0.1) for each population detected by our method, the ratio of PRDM9-A/C binding motifs in the African population (PRDM9- A ratio: 50.4%; PRDM9-C ratio: 33.3%) is much higher than that of the other populations (Fig. 5B). On the other hand, as we define enrichment factor by considering the recombination rate of the entire region around the motif, the enrichment factor of PRDM9- C binding motifs in the African population is corrected to be on the same level as the other populations due to the higher overall recombination rate in the population (Fig. 5C). Although using the absolute value of the recombination rate to show the contribution of recombination hotspot determinants is straightforward, our framework provides an alternative quantification method by utilizing the relative criterion, which may be more robust to the local region noise and population batch effect. The relation between the recombination rate and the enrichment factor of a motif is complex, which cannot be modeled with a linear regression (Fig. 5D-E), with the Pearson correlation coefficient being -0.034 and the *R*^2^ score being 1.79 ∗ 10^−3^ on motifs extracted from 5 populations. However, together with the recombination rate, by quantifying the relative contribution, our method provides insights into the recombination hotspot determinants.

**Fig. 5.**
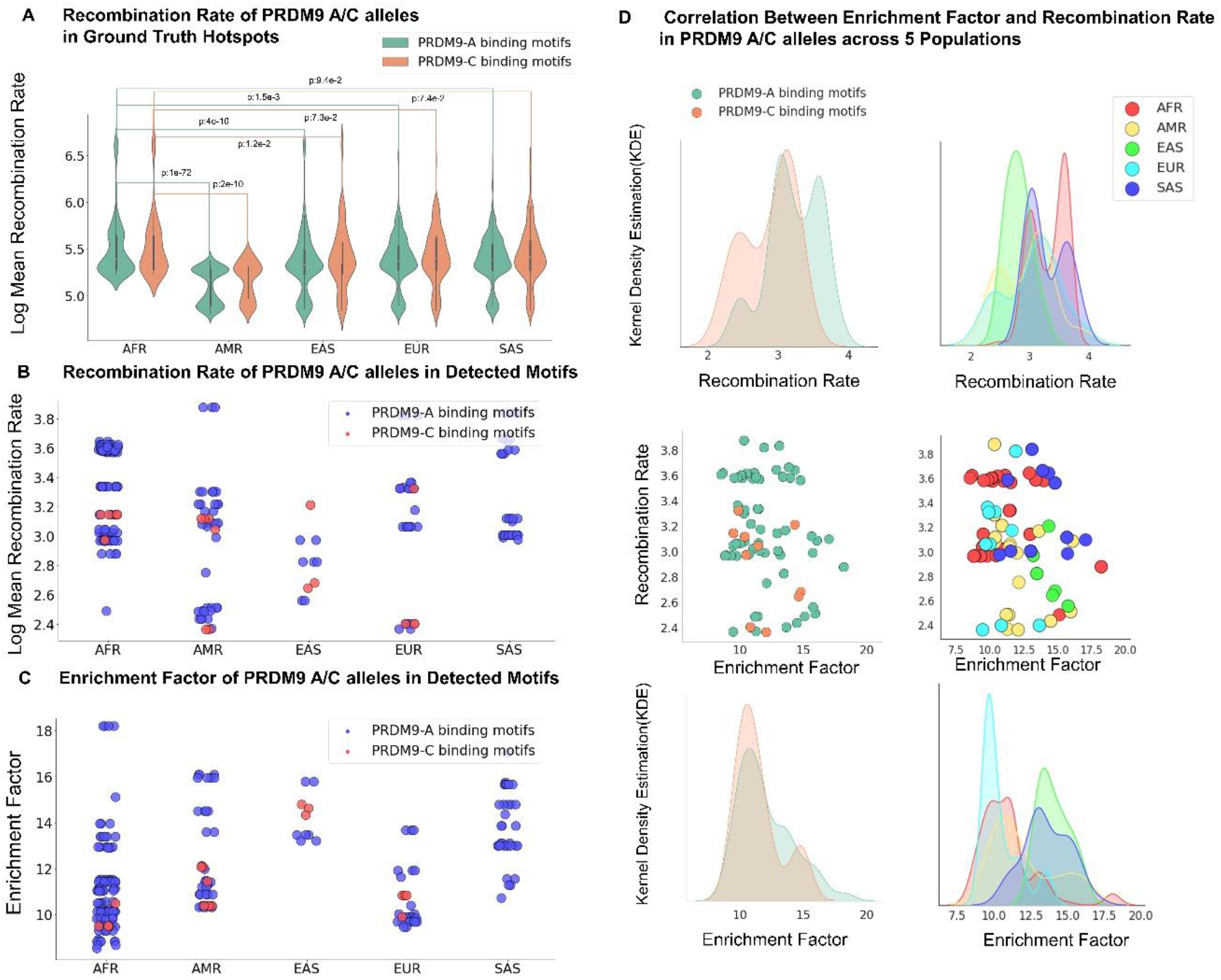
Recombination rates and enrichment factors of PRDM9 alleles within different populations. **(A)** Recombination rates of ground truth hotspots at different PRDM9 binding motifs in five populations, normalized by log average recombination rate. The African population has significantly higher recombination rates in both alleles compared to the other populations. **(B)** Recombination rates of the RHSNet-identified PRDM9-A and PRDM9-C binding motifs. The recombination rate of PRDM9-A/C alleles in the African population is significantly higher than that in other populations. **(C)** Enrichment factors of the RHSNet-identified PRDM9-A and PRDM9-C binding motifs. The enrichment factor of PRDM9-C binding motifs in the African population is corrected to be on the same level as the other populations due to the higher overall recombination rate in the population. **(D)** Paired relation between enrichment factors and recombination rates among all the detected PRDM9-A/C alleles across different populations. We show the recombination rate distribution of all the PRDM9-A/C alleles together and within each population in the upper row. As shown in the middle row, the relation between enrichment factors and recombination rates is more complex than linear correlation. In the bottom row, we illustrate the enrichment factor distribution of all the PRDM9-A/C alleles together and within each population.

### Generalization to PRDM9-lacking species and sensitivity to sex differences

The recombination event has been studied in a broad range of species(*11, 34–36*), including humans, mice, yeast, birds, and pigs. In addition to the human data, our method can be further applied to other species, regardless of having the PRDM9 gene. Although PRDM9 is shown to be the major determinant of recombination hotspots in both humans and mice, the predicted binding motifs are different in the two species(*4*). We apply our method to the Mice data(*11*) and identify the most significant determinant motifs. In addition to the GT-rich motifs, which are enriched in the SPO11-oligo hotspots and the usual mice PRDM9 binding sites(*11*), surprisingly, we also identify an AC-rich motif (Fig. S9A). Although this motif has not been studied extensively in the mice-related literature, it is reported as a part of the binding motif for the PRDM9^9R^ zinc finger binding domain(*11, 29*). Unlike apes and mice, birds and yeast lack a PRDM9 gene, leading to different recombination hotspot patterns in these species(*34–36*). On the Yeast dataset(*6*), the poly-(A) motif is identified as the most significant determinant in hotspots (Fig. S9A), which is completely different from mice and humans. However, the result is consistent with the previous study, which demonstrates that Poly-(A) motif occurs more frequently in the hotspots(*6*). Moreover, the motifs enriched in the yeast promoters(*37*) are also predicted to be of vital importance to the recombination event (Fig. S9A), which supports the theory that, in the PRDM9-lacking species, hotspots are highly conserved due to the natural selection pressure(*34*).

The recombination differs between males and females of the same species, including humans and mice(*17, 38*). The females show a higher overall recombination rate and more complex crossovers than the males(*2, 12, 16*), despite the elusive mechanisms behind the differences(*2*). Although most of the recombination hotspots (>88%) are shared between males and females, the strongest hotspots tend to be sex-specific(*17*). In our study, the most significant motifs detected from the Icelandic males are conserved PRMD9 binding motifs (Fig. S9B), which is consistent with the finding that the PRDM9-binding sites are frequently methylated at male-biased hotspots(*17*), although the motifs can be different in different species. On the other hand, the contributing determinant motifs of the Icelandic females are less conservative (enrichment factor: 8.76 ± 1.31) than the paternal ones (enrichment factor: 15.31 ± 2.01). Also, in the female, the determinant motifs are much more diverse than those in males, which may result from the distinct methylation mechanism (unlike males, DNA methylation increases in the region ±75bp adjacent to the PRDM9-binding sites), more complex crossover, and higher evolution speed(*1, 2, 17*) (Fig. 3C, female rate: 52.48 ± 67.29 *cM/mb*; male rate: 39.53 ± 40.63 *cM/mb*). Although the data themselves may not be sufficient to illustrate the mechanism behind the sex biases in recombination, the identified and quantified determinants suggest that, in females, diverse factors, including PRDM9 and SPO11 (the rank 2 motifs), control the female-biased hotspots, while, in males, the hotspots tend to be PRDM9-directed.

### Recombination hotspot motif embedding for evolutionary determinants discovery

We extend our method to identify and quantify the recombination hotspot determinants more intuitively and systematically. Instead of only listing the detected determinants and their enrichment factors, we learn the representation of the motifs in a 2D space, visualizing and clustering them in such a space. To avoid black-box modeling, we also visualize the physical meaning of the motifs with heatmaps (Fig. 6). Within the Icelandic dataset (Fig. 6A), the standard deviation of the determinant motif embeddings in the maternal population is much larger than that in the paternal population (maternal: 0.0447 ± 1.9 ∗ 10^−2^, *p* = 3.11 ∗ 10^−6^; paternal: 0.0355 ± 1.6 ∗ 10^−2^; Materials and Methods), which suggests that, in females, the recombination hotspot determinants are more diverse and less conservative than that in the males. This finding further supports our hypothesis that diverse factors contribute to the female-biased hotspots (Section 5).

**Fig. 6.**
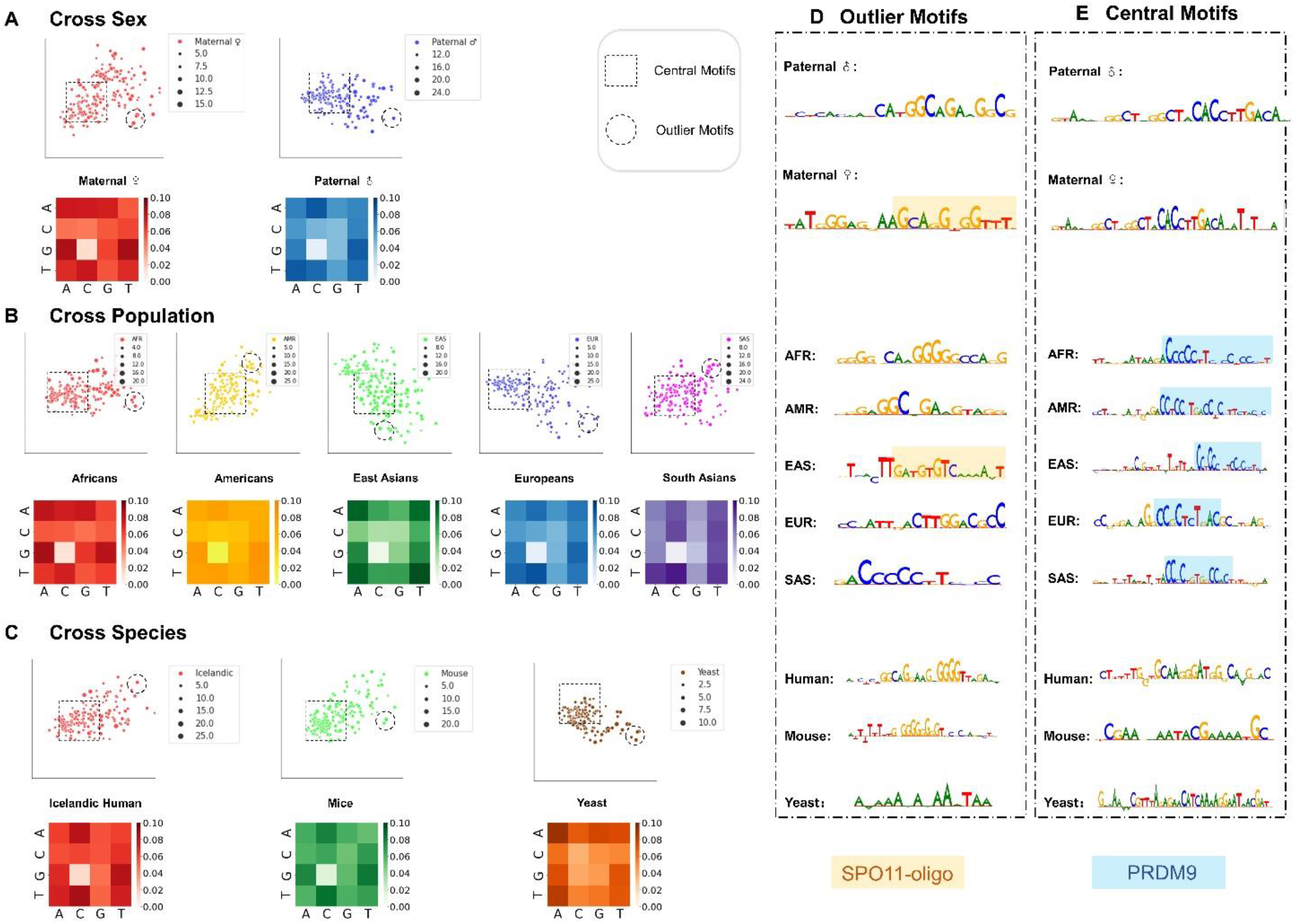
Motif embedding and outlier detection for recombination hotspot determinants discovery. **(A)** Visualization of the motif embeddings over different sexes in the Icelandic dataset. The size of the scatter reflects the enrichment factor of that motif. Each heatmap below the cluster suggests the 2-mer appearance frequency in the detected motifs. The representative central and outlier motifs are shown in **(D)** and **(E)**. **(B)** Visualization of the motif embeddings across different populations. The enrichment factors differ significantly between central motifs and outlier motifs across populations. **(C)** Visualization of the motif embeddings across different species. Poly-(A) motifs are the most enriched outlier motifs in Yeast hotspots. Also, the Human/Mice outlier shows evolutionary discovery in motifs correlated with SPO11 oligos.

For all the visualizations of the motifs across different sexes (Fig. 6A), different populations (Fig. 6B), and different species (Fig. 6C), the motifs within the central area of the embedding space tend to have a smaller enrichment factor value, represented by the size of the point, than the outlier motifs. Because the enrichment factor value shows the importance of the determinant, investigating the outlier motifs may identify the evolutionary important motifs of the population and species. In the Icelandic population, within the central region of the embeddings, the motifs in both the maternal population and the paternal population are PRDM9-related motifs (Fig. 6E). However, regarding the outliner motifs, they are very different. For males, the motifs tend to be short and strong, while the motifs are long and diverse in the females. Similar to the Icelandic data, within the 1000 Genomes Project dataset, the enrichment factor differences between central motifs and the outliers across different populations share a similar pattern (Fig. 6B), where enrichment factors differ significantly between central motifs and outlier motifs across populations, especially in East Asians (AFR: *p* = 2.85 ∗ 10^−4^ ; AMR : *p* = 8.51 ∗ 10^−2^ ; EAS: *p* = 1.12 ∗ 10^−5^; *EUR*: *p* = 1.16 ∗ 10^−2^; SAS ∶ *p* = 0.23 ). For the convenience of the study, we randomly select the most distinct outlier motifs in different populations across the embedding space and visualize them in Fig. 6D. Interestingly, the motifs enriched in the SPO11-oligo hotspots(*11*) show up in the East Asian population. Although the molecular studies are mainly performed on Mice(*11*), and researchers have not performed such studies on different human populations systematically, our method provides the first quantitative depiction of the recombination hotspot determinant motifs across diverse populations.

We further extend our analysis to different species (Fig. 6C). A similar pattern appears. The central motifs, shared by different species, have smaller enrichment factors than the outlier motifs, which are likely to be species-specific (Icelandic human: *p* = 1.57 ∗ 10^−25^; Mice: *p* = 2.2 ∗ 10^−8^; Yeast: *p* = 7.4 ∗ 10^−5^). For example, the poly-(A) motifs are the most important ones for yeast, which does not have the PRDM9 gene(*6*). On the other hand, our method provides a new way to define the evolutionary distance between different species(*39*), using the embedding of the recombination hotspot determinants. By calculating the difference between two embedding distributions, we quantify the difference between two species via Maximum Mean Discrepancy(*40*) (MMD) (Materials and Methods). For example, using our method, the evolutionary distance between Human and Mice (0.0221, *p* = 0.9774) is much smaller than that between Human and Yeast (0.3729, *p* = 8.5 ∗ 10^−3^).

## Discussion

Recombination is one of the most important processes in miosis for sexually reproducing organisms, which can produce genetic diversity for natural selection. Despite its important role in evolution, people know little about the entire process and its molecular mechanism. Although a large amount of data have been accumulated from various giant projects, such as HapMap(*28*) (3.1 million human single nucleotide polymorphisms (SNPs) genotyped in 270 individuals), Sperm(*13*) (31,228 human gametes from 20 sperm donors), and 1000 Genomes Project(*18*) (84.7 million SNPs of 2,504 individuals), seldom have researchers developed methods to analyze data from different studies and even different species.

Here, we propose a new computational method, RHSNet, which enjoys the strength of deep learning(*24*), activation backpropagation(*26*), and signal processing(*27*), to identify and quantify the recombination hotspot determinants. Although our method is not designed specifically for recombination hotspot region prediction, it can outperform almost all the previous methods in this task across different studies, different populations, different sexes, and different species. More importantly, RHSNet can identify and quantify the determinants that contribute significantly to the recombination hotspot formation. In addition to quantifying the relation between PRDM9 binding motif(*4, 8–10*), histone modification(*3, 11*), GC content(*2, 12*) and recombination hotspots, it reveals the contribution of different PRDM9 alleles in different populations. Further studies on different species, including PRDM9-lacking species, and different sexes suggest the generalization power and sensitivity of the proposed method. The cross-sex, cross-population, and cross-species studies show the potential of our method to identify the evolutionary determinants. Although RHSNet is purely data-driven and more work can be done to further improve it, including using the gene annotation related to the location information(*41*), chromatin accessibility information(*42*) as well as the conditional analysis(*43*), it is potentially helpful to assist researchers in illuminating the mechanisms underlying recombination and evolution.

## Materials and Methods

### Dataset construction

In our study, we use datasets from a number of projects, including the Icelandic(*2*) dataset, the HapMap II(*28*) dataset, the Sperm(*13*) dataset, the 1000 Genomes Project(*18*) dataset, the Mice(*11*) dataset, and the Yeast(*6*) dataset.

The Icelandic(*2*) dataset, provided by 1,476,140 crossovers from 56,321 paternal meiosis and 3,055,395 crossovers from 70,086 maternal meiosis, has a 642 bp resolution (655 bp for the paternal part) generated from Icelandic pedigrees on the GRCh38 human reference genome(*44*), from which we select 20,000 hotspots with an average recombination rate of 51.07 cM/Mb and 20,000 coldspots with an average recombination rate of 1.78 ∗ *e*^−10^ cM/Mb (resolution from 500 bp to 1000 bp) for cross-validation. Based on the fact that, in the sex-average map, the average length of those hotspots is relatively shorter (averaging 526 bps) than that of coldspots (averaging 3,071 bps), we sort the recombination rates of all the possible sequences and select the lowest 20,000 coldspots with a proper resolution to construct the negative samples.

The HapMap II(*28*) dataset is segmented into hotspots (average rate 10.5 cM/Mb) and coldspots (average rate below 0.5 cM/Mb) regions based on a hidden Markov model with emission probabilities defined as *p*(*observedrate*|*hot*) and *p*(*observedrate*|*nonhot*). Hotspots longer than 4kb are discarded for localized recombination events, and the matched set of coldspots are defined through a greedy searching method for a 300kb region with sequence having GC content matching.

The Sperm(*13*) dataset is built with 31,228 sperm cells from 20 sperm donors, among which 813,122 crossovers from 787 aneuploid chromosomes are identified. The recombination rates vary in 20 sperm donors ranging from 22.2 to 28.1 crossovers per cell. A fine-scale genetic map is generated by us from those 813,122 crossovers events (Data and materials availability) by stepping along the genome at 500kb intervals, dividing the number of crossovers that occurs up to each point and by the total number of cells. We further select 5,000 hotspots with an average recombination rate of 19.96 cM/Mb and 5,000 coldspots with an average recombination rate of 1 ∗ *e*^−20^ cM/Mb for cross-validation.

The Mice dataset(*11*) is constructed according to the SPO11-oligo maps from C57BL/6J. Soluble protein is subjected to two successive rounds of affinity purification with a monoclonal anti-mSPO11 antibody and protein A-agarose beads. Sampled hotspots are identified with 0.000377 RPM/bp, which is 50 times of the average reads per million (RPM) within the mappable GRCm38/mm10 genome. The selected hotspots are further cropped into 1000bp for deep learning purposes, resulting in 9,620 hotspot sequences. Similarly, the coldspot region is equally selected with 1 ∗ 10^−7^ times of the average reads per million (RPM) within the mappable GRCm38/mm10 genome region and cropped with the same length.

In order to verify our method’s generalization ability on species lacking the PRDM9 gene, we further construct Yeast(*6*) (Saccharomyces cerevisiae) hotspots from nearly 52,000 markers in all the four viable spores derived from 51 meioses of an S288c/YJM789 hybrid strain. Pairs of genotype changes isolated from all other changes are called NCOs if they appear on the same spore, or COs if they appear on two spores. In total, 468 meiotic hotspots (averaging 842bp in length) that contain 92 COs and 74 NCO are cropped from the S288C Yeast reference genome. As S288C Yeast reference genome is much shorter than that of humans and mice, we define the corresponding recombination coldspots as the gap sequences between two recombination hotspots with at least 1000bp away from hotspots.

We utilize the population-wise recombination maps generated from the 1000 Genomes Dataset(*18*) to conduct the direct meiotic recombination hotspot prediction as well as downstream analysis. Fine-scale maps having an average resolution of 711 bp based on 26 diverse human populations are further merged into five super-populations: African (AFR), admixed American (AMR), East Asian (EAS), European (EUR), and South Asian (SAS). The merged five super-populations share a high Spearman correlation ranging from *ρ* = 0.986 to *ρ* = 0.998 within each category. Therefore, we further map the recombination hotspot regions within each super-population category to the GRCh38 reference genome, and generate corresponding hotspots for each population (AFR 50,049 hotspots avg 27.46cM/ Mb; AMR 18,160 hotspots avg 27.32cM/Mb; EAS 27,030 hotspots avg 38.44cM/ Mb; EUR 31,283 hotspots avg 35.05cM/ Mb; SAS 32,593 hotspots avg 33.92cM/ Mb). Similar to the Icelandic(*2*) dataset, we select the coldspot sequences from the lowest recombination rate regions of the recombination map (AFR 50,049 coldspots avg 0.0378cM/Mb; AMR 18,152 coldspots avg 0.039cM/Mb; EAS 27,020 coldspots avg 0.094cM/Mb; EUR 31,273 coldspots avg 0.038cM/Mb; SAS 32,583 coldspots avg 0.033cM/Mb). Statistical comparison between the generated hotspots and coldspots data can be found in Table S3.

When targeting sex-specific recombination prediction, we use the ChIP-seq features as extra information for the proposed RHSNet. Following the previous research(*2*), we use histone modifications from ovary for the maternal map and testis for the paternal map. On the Icelandic(*2*) dataset, we define the hotspot ChIP-seq feature as the closest narrow peak next to the hotspot sequence found in six different kinds of histone modifications(*45*) (H3K4me1(*31*), H3K4me3, H3K27ac(*32*), H3K9me3, H3K36me3, H3K27me3). Similarly, we define the coldspot feature as the closest narrow peak next to the coldspot sequence. Specifically, the feature vector is set to zero when the actual distance exceeds 10kbp. The ovary histone modifications are downloaded from the ENCODE portal(*46*) with the following identifiers: **ENCSR139TLA, ENCSR268JQE, ENCSR113AFY, ENCSR659MYS, ENCSR956UFV, ENCSR037SNV**. The testis histone modifications could be found by following identifiers: **ENCSR611DJQ, ENCSR136ZQZ, ENCSR956VQB, ENCSR561MYM, ENCSR376JOC, ENCSR503QSX.**

### Deep learning network architecture

The deep learning network architecture consists of two independent feature extractors. The sequence feature extractor first encodes DNA sequences into one-hot matrices. To represent each nucleotide, we define the encoding as a vector of size 4, A as (1 0 0 0), T as (0 0 0 1), G as (0 0 1 0), and C as (0 1 0 0). Next, two convolution layers are introduced as a feature extractor to encode the one-hot matrix into a relatively shorter feature vector. Utilizing sequential neural networks known as Gated Recurrent Units (GRU) and multi-head Attention mechanism, we take advantage of the combination of sequential model and attention model to capture the deep contextual information of the input sequence.

The ChIP-seq feature extractor takes the sequence’s nearest corresponding ChIP-seq(*47*) information (including the peak value, score, and signal value from 6 different histone modifications: H3K4me1, H3K4me3, H3K27ac, H3K9me3, H3K36me3, and H3K27me3) as input. The ChIP-seq information is encoded as high-dimensional feature vectors and fed into the network. We apply fully connected layers with dropout at this part of the network to prevent over-fitting during training. Finally, high-dimensional features containing both sequences and their surrounding histone H3 protein information are passed to the final SoftMax layer, producing the final prediction. Detailed illustration of our prediction model can be referred to Fig. S2.

### Parameter settings and implementation details

The baseline CNN has two plain convolution layers connected with the final SoftMax layer. The first 1D convolution layer is designed with a filter of length16 and a kernel size of 30 with ReLU activation. The proposed RHSNet connects a Gated Recurrent Units (GRU) and a Multi-head attention layer after the 2-layer feature extractor. The number of neurons assigned in GRU is designed as 16. We use four multi-heads with the head size as 4 in the multi-head Attention mechanism.

Stochastic gradient descent with the momentum parameter as 0.9 and dynamically updated weight decay is used to optimize the learning process. The batch size is selected to be 64. The initial learning rate is set to be 1 ∗ *e*^−3^. The dropout ratio is set to be 0.1 after the first and the second convolution layer to prevent over-fitting.

### Low-pass filter-based signal extraction from guided backpropagation

The previous method, known as DeepLIFT^26^ (Deep Learning Important Features), successfully assigns contribution scores of input DNA sequences. By calculating the gradient of each activated neuron through guided backpropagation, contribution scores can be computed efficiently in a single backward pass based on the reference sequence with (A, T, C, G) distributed with probabilities as 0.3, 0.2, 0.2, and 0.3, respectively. With the reference sequence 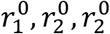 as input, the reference activation *y*^0^ could be computed as:

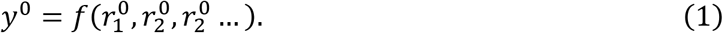

The key idea of important score extraction is through guided backpropagation. For each neuron *y*, Δ*y*^+^ and Δ*y*^−^ are defined as having positive and negative component of Δ*y*:

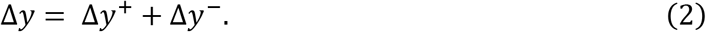

Now, given an input neuron *x* and the target neuron *t*, there is a difference of Δ*t* from the reference neuron *r*. The multiplier *m*_Δ*x*Δ*t*_ could be defined as the contribution of Δ*x* to Δ*t* divided by Δ*x*:

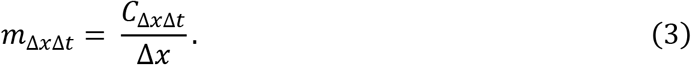

Also, given *C*_Δ*xi*Δ*yi*_ along with *C*_Δ*yi*Δ*t*_, we can show that the definition of *m*_Δ*x*Δ*t*_ according to the chain rule would satisfy summation-to-delta:

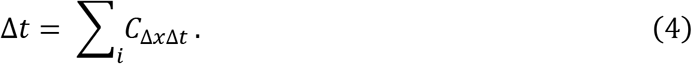

Notably, using the chain rule in which the input layers have one-hot encoded sequence *s*_1_, *s*_2_, … *s*_*n*_, hidden layers *y*_1_, *y*_2_, … *y*_*n*_, and the target output *t*, based on equation (**4**), we can have:

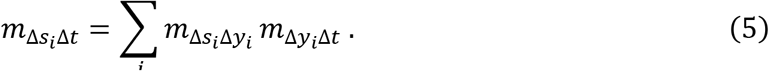

We can compute the multipliers for each input sequence *s*_*i*_ efficiently via backpropagation.

Inspired by DeepLIFT, a more advanced TF-MoDISco(*48*) introduced for transcription factor prediction was proposed. However, DeepLIFT and TF-MoDISco share a common disadvantage of having a strong assumption on the discovered motif length based on previously known probabilistic motif models. Such a strong assumption is further enhanced when adjusting the sliding window size similar to the expected length of the core motif and its flanks, which can be unknown for innovative motif discovery. Finding longer motifs is crucial in the recombination hotspot prediction task because each PRDM9 zinc finger is 28 amino acids long and is usually decoded within an 84 bp repeating tandem(*4*). Also, the PRDM9c (ZF8–13) motifs are usually 21bp long, because they are accompanied by 5’ (five prime) and 3’ (three prime), making the traditional sliding-window-based method less efficient.

Instead of simply providing a sliding window with a certain kind of size, which gives a strong assumption on the discovered motif length, RHSNet offers a general solution as a localization method for innovative motif discovery with variant lengths. In our motif extraction algorithm, we utilize low-pass filters with different kinds of factors and a peak detection algorithm, from which we could easily control the length range of the detected motifs by controlling the low-pass factor.

The contribution scores generated from backpropagation are firstly transformed into one-dimensional signals and fed into the low-pass filter. Opposite from high-pass filters, the low-pass filter allows low-frequency signals to pass:

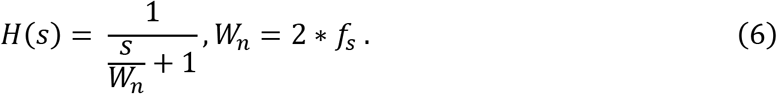

The order of the proposed low-pass filter is fixed as 8, and the frequency scalar is flexibly chosen from [0.1,0.2,0.4]. Therefore, when selecting frequency scalar as 0.4, *W*_*h*_ = 0.4 = 2 ∗ 200/1000, we have the Nyquist frequency as *f*_*s*_ = 200Hz. Similarly, when selecting frequency scalar as 0.1, *W*_*n*_ = 0.1 = 2 ∗ 50/1000, we have the Nyquist frequency as *f*_*s*_ = 50Hz.

The low-pass filter provides a smooth form of signals by eliminating short-term fluctuations and retaining long-term development trends, in which longer motifs enriched by a relatively high-frequency signal are reserved. By detecting each peak with its nearby valley, we could easily extract the motif in the middle. We choose 0.06 as the prominence parameter for the peaks. Also, the valleys are defined as the peaks of the reversed signal where the interval of each valley width is set to 1.

### Enrichment factor definition

The motif enrichment factor is defined by the ratio of the selected motif’s contribution score over the average contribution score of the entire input sequence through backpropagation. Different from the recombination rate, which is an absolute value, the enrichment factor is more of a relative index which indicates how strong the enriched motif signal is among the entire input sequence. That is, the larger the enrichment factor, the higher chance that such a cropped motif plays a more important role in the recombination events.

### ChIP-seq feature extraction and importance score board

We quantify the distribution of three Chromatin Immunoprecipitation Sequencing (ChIP-seq) features (Score, Signal Value, Peak Value) extracted from the peaks of signal enrichment based on six different kinds of histone modifications(*45*) (H3K4me1(*31*), H3K4me3, H3K27ac(*32*), H3K9me3, H3K36me3, H3K27me3), on different cell lines. Specifically, for maternal recombination map, we select the histone modifications from homo sapiens ovary tissue (female adults). For maternal recombination map, we select the histone modifications from homo sapiens testis tissue (male adults). Score is an integer value ranging from 0 to 1000, representing the significant score of each peak. Signal Value measures the average enrichment for the related peak region. Peak Value is the point-source called for this peak. It is the 0-based offset from the chromosome starting point and is set to -1 if no point-source is called. We take the log mean feature values of each histone modification and choose the nearest conservative peaks of each hotspot/coldspot clip.

Within each type of histone modification, p-values are obtained using the two-tailed Student’s t-test. The calculated t-statistics of H3K4m3-signalValue (8.03 ∗ 10^−7^), H3K4m3-peakValue (1.16 ∗ 10^−9^), and H3K36me3-peakValue (3.91 ∗ 10^−2^) show the significant statistical difference between hotspots and coldspots within 18 features.

To further quantitively perceive the difference between the above 18 features and provide a more intuitive impression of their significance, we define the importance scorecard (Fig. 4E) not only as an index for measuring the usefulness of each feature, but also as an important index for measuring the quality and statistical significance of the feature’s contribution to the improvement of the final prediction. The score is calculated by backpropagating the activation through the entire network to the ChIP-seq feature extraction branch of the RHSNet deep learning model. The greater the contribution score, the higher likelihood that this feature, along with its histone modification, plays a critical role in the prediction process.

### PRDM9-A/C allele identification

All the PRDM9-A/C alleles are identified through rigorous multiple sequence alignment from both ground-truth hotspots and detected binding motifs. The major PRDM9-A allele: *CCNCCNTNNCCNC* and its reverse: *GGNGGNANNGGNG*, as well as PRDM9-C allele: *CCGCNGTNNNCGT* and its reverse: *GGCGNCANNNGCA*, are selected as the reference sequences. Each discovered motif would be conducted a pairwise sequence alignment with each allele using a dynamic programming algorithm. The selected motif would only be considered containing the major allele A when having the minimum alignment score of 8, which is chosen as the certainty of the 8 certain bases: *CC-CC-T--CC-C* and *GG-GG-A--GG-G*. The identification rule is also applied for the identification of the rare allele C.

### Motif embedding and outlier detection

To intuitively visualize the discovered motif, we utilize DNABERT(*49*), which extracts the short- to long-term patterns of each enriched motif into a fixed-size embedding vector. The embedded vector is further fitted into t-distributed Stochastic Neighbor Embedding(*50*) (t-SNE) to visualize the 2-dimensional embedding vector and investigate their divergence across different sexes, populations, and species. Each heatmap under Fig. 6A.B.C illustrates the physical meaning of each motif embedding cluster. Practically, we calculate the frequency of the 16 possible 2-mers within the [A, T, C, G] alphabets that appear in each population, sex and species, and visualize them with the saliency heatmap.

The outliers within each cluster are defined by the Local Outlier Factor (LOF)(*51*). This algorithm is an unsupervised anomaly detection method that computes the local density deviation of a given data point with respect to its neighbors. It considers as outliers the samples that have a substantially lower density than their neighbors. During the outlier detection, we compute the locality based on the given k-nearest neighbors, whose distance is used to estimate the local density. By comparing the local density of a sample to the local densities of its 50 neighbors, we can identify samples that have a substantially lower density than their neighbors. These motifs are considered outlier motifs.

Regarding calculating the divergence of the embedded vectors distributing within 2-D space, we calculate the average distance of each embedded vector of selected species/sex/population to the central point of each embedding space. The maternal population has an average distance of 0.0447 (±1.9 ∗ 10^−2^), which is much larger and has grater dispersion than that of paternal crossovers: 0.0355 ± 1.6 ∗ 10^−2^.

### Maximum Mean Discrepancy (MMD) calculation

As a kernel-based distance calculation metric, Maximum Mean Discrepancy (MMD) can accurately quantify the similarity and the distance between two vector distributions. When measuring the difference between two embedding distributions, because we need to measure the non-parametric distribution distance between the source and target data, namely, P and Q, we use MMD. The calculation is done by the following equation:

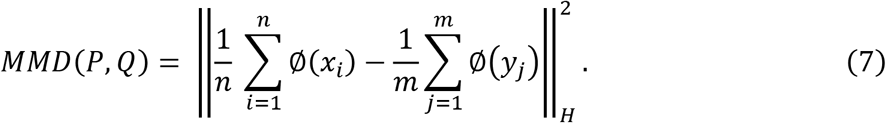

For example, when comparing the embedding distance between paternal motif vectors (P) and maternal motif vectors (Q), we map P and Q (Original *X* space) from the original embedding space to another space H (Hilbert space) through the function ∅: *X* → *H*. Then, we can calculate the mean difference between P and Q in the H space feature dimension. When utilizing MMD as the evaluation metric for calculating the distance between two distributions within the embedding space, we can determine whether the two distributions are similar. Quantitatively, the *MMD*(*Human*, *Mouse*) is 0.0221 (*p* = 0.9774), which is much smaller than *MMD*(*Human*, *yeast*)=0.3729 (*p* = 8.5 ∗ 10^−3^).

## Acknowledgments

**Fig. 1** is created by Heno Hwang, a scientific illustrator at King Abdullah University of Science and Technology (KAUST).

## Author contributions

Conceptualization: Y.L, X.G

Methodology: Y.L, S.C, T.R

Investigation: Y.L, S.C

Visualization: S.C

Supervision: Y.L, X.G

Writing—original draft: Y.L, S.C

Writing—review & editing: Y.L, X.G, K.Y.Y, H.K.

## Data and materials availability

The RHSNet source code, the motif extraction script, and the pretrained models are available on GitHub under the MIT License: https://github.com/frankchen121212/RHSNet.

Recombination hotspots and coldspots datasets for training and testing are acquired from multiple data sources across different studies, sexes, populations, and different species, and are now made available at: https://drive.google.com/drive/folders/1kNMrZgsceA_ZuwhEDjEeiA3eSBHix2Cv?usp=sharing.

The genetic maps for Icelandic(*2*) dataset accompanied by its paternal and maternal maps are obtained from: https://science.sciencemag.org/content/363/6425/eaau1043/tab-figures-data.

Our generated genetic maps for Sperm(*13*) dataset along with the defined hotspots are created from : https://zenodo.org/record/3561081#.YK5FY6gzYYo.

The HapMap II(*28*) dataset together with the source code for Equivariant CNN model are obtained from: https://github.com/luntergroup/EquivariantNetworks.

Recombination maps of 26 populations as well as corresponding five super-populations: African (AFR), admixed American (AMR), East Asian (EAS), European (EUR), and South Asian (SAS) of the 1000 Genomes Project(*18*) on hg38/GRCh38 are available at: https://drive.google.com/drive/folders/1Tgt_7GsDO0-o02vcYSfwqHFd3JNF6R06. The Yeast hotspot data are available via: https://pubmed.ncbi.nlm.nih.gov/18615017/

## Supplementary Materials

### Evaluation Criteria

We use multiple evaluation methods to quantify the prediction results generated by different prediction methods according to the number of TP (True Positives), FP (False Positives), FN (False Negatives), and TN (True Negatives) samples.

Accuracy (ACC) can judge the performance of our model, but there is a serious flaw: in the case of imbalanced positive and negative samples, the category with a large proportion will often become the most important factor affecting accuracy. Therefore, sometimes, it might not reflect the overall prediction performance of the model. Accuracy is defined as follows:

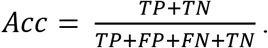

F1 score (F1-Score) is a weighted average of recall and precision, and is defined as follows:

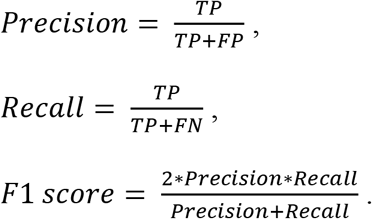

Matthews correlation coefficient (MCC) is an index used in machine learning to measure the binary classification performance of the predictor. It is generally considered to be a relatively balanced evaluation metric, and it can be applied even when the number of positive and negative classes is extremely imbalanced. MCC is essentially a coefficient describing the correlation between the actual classification and the predicted classification. Its value range is [-1,1]. A value of 1 indicates a perfect prediction of the subject, and a value of 0 indicates that the predicted result is not as good as the random predicted result. -1 means that the predicted classification is exactly the opposite of the actual classification. MCC is defined as:

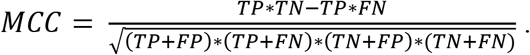

ROC curve and the corresponding AUROC score are other evaluation indexes. The larger the area under the curve (AUC), or the curve closer to the upper left corner (true positive rate=1, false-positive rate=0), the better the model’s prediction in the task.

### Imbalanced Testing

The recombination hotspots prediction is a problem that the number of positive samples (hotspots) is much less than that of negative samples (coldspots), which makes it difficult for the predictor to achieve high sensitivity. In average maps of the Icelandic2019(*53*) dataset, we found 130,172 hot spots and 755,041 cold spots with extreme diverse recombination rate from1.7e10−17cm/Mb to 56,242.73cm/Mb. Those sequences with lengths longer than 1kb were discarded during training, and the remaining hotspot samples are 17 times less than the cold spots samples (see **Table S1**).

As shown in **Fig. S4**, after the balanced training, we test our model throughout all the sequence samples across the entire genome from the paternal and maternal maps at the Icelandic 2019(*53*) dataset. The area under the receiver operating characteristic (AUROC) was calculated to evaluate the algorithm’s performance under an imbalanced prediction task. The RHSNet approach achieves the best AUCROC score of 0.689.

### Recombination Rate Comparison

The illustration of the recombination rate distribution over the detected paternal and maternal motifs (**Fig. S14**) shows that the maternal recombination rates are relatively higher than that in paternal crossovers within each rate interval.

### Hit@20,50,100 Evaluation

Similar to the Recommendation System (RS) ranking evaluation index, the motif detection method proposed in RHSNet that recommends prediction motifs could be evaluated similarly.

First, we calculate the enrichment factor for each detected motif through the contribution score of that slot over the entire input sequence. In this way, each detected motif will get an enrichment factor score. Furthermore, we sort these scores in descending order so that the motifs with the highest enrichment factor would be ranked in the front.

According to the above ranking function, we can count whether the PRDM9-A/C allele exists for each detected motif is in the top 20 of the sequence, and if so, we add one count to Hit@20. In the end, the top 20 number/total is Hit@20. Similarly, Hit@50 and Hit@100 are the top 50/100 detected PRDM9-A/C alleles over the total number of motifs. Furthermore, the value of Hit@20 may exceed 20 because the detected motif is usually 21bp long, and it might contain more than one 12-bp motifs in one sequence.

Intuitively speaking, the key factor of predicting an input sequence as the recombination hotspot will give much credit to the PRDM9-A/C allele. As showed in **Table S6,** the PRDM9-A allele is ranked pretty high in Hit@10/20/100 evaluation, demonstrating that RHSNet could precisely identify the key factor of the recombination hotspot determinant. Also, the number of the RHSNet-detected PRDM9-A allele is approximately nine times larger than that of the PRDM9-C allele. Such a result is consistent with previous studies that the PRDM9-A motif plays a role in approximately 40% of hotspots and is proposed to be involved in initiation specification or other aspects of recombination activity.

---

**Fig. S1.**
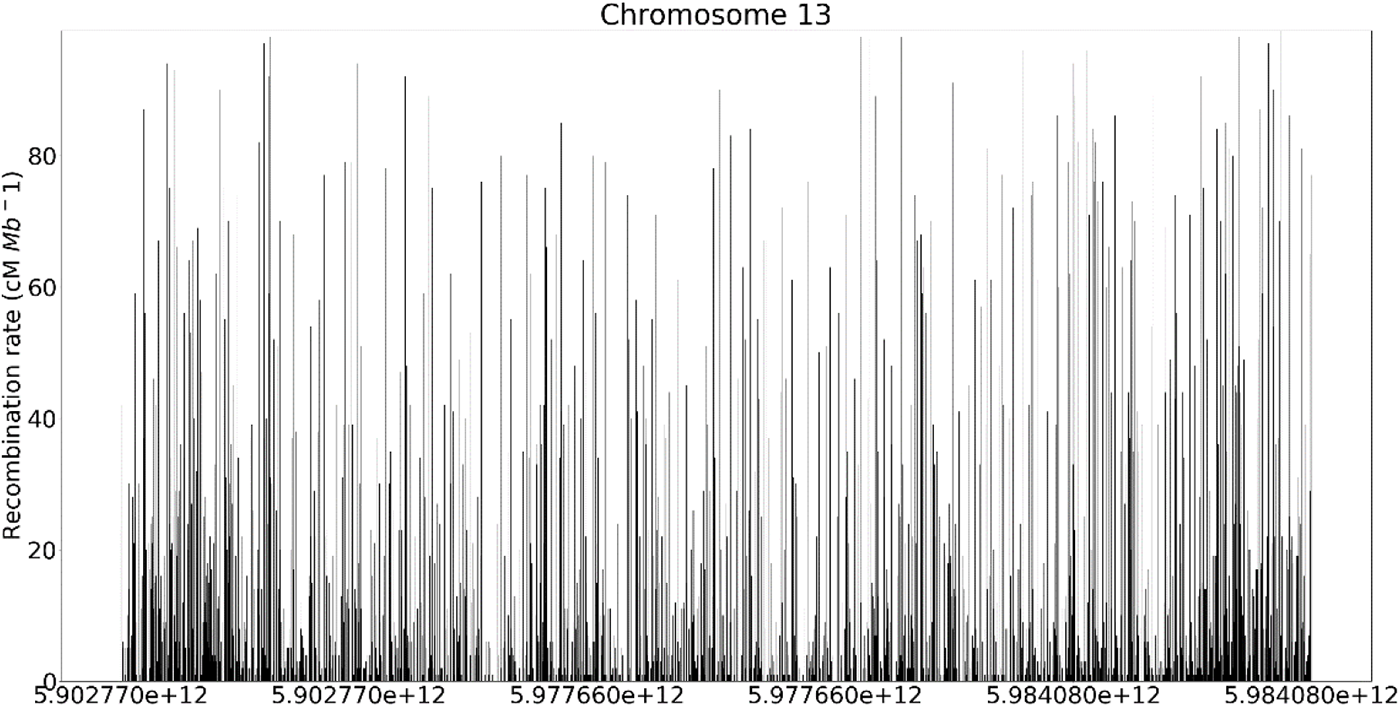
Recombination rate distribution over chromosome 13. The recombination rate distribution over chromosome 13 from the Icelandic dataset. The resolution is set as 100kbp.

**Fig. S2.**
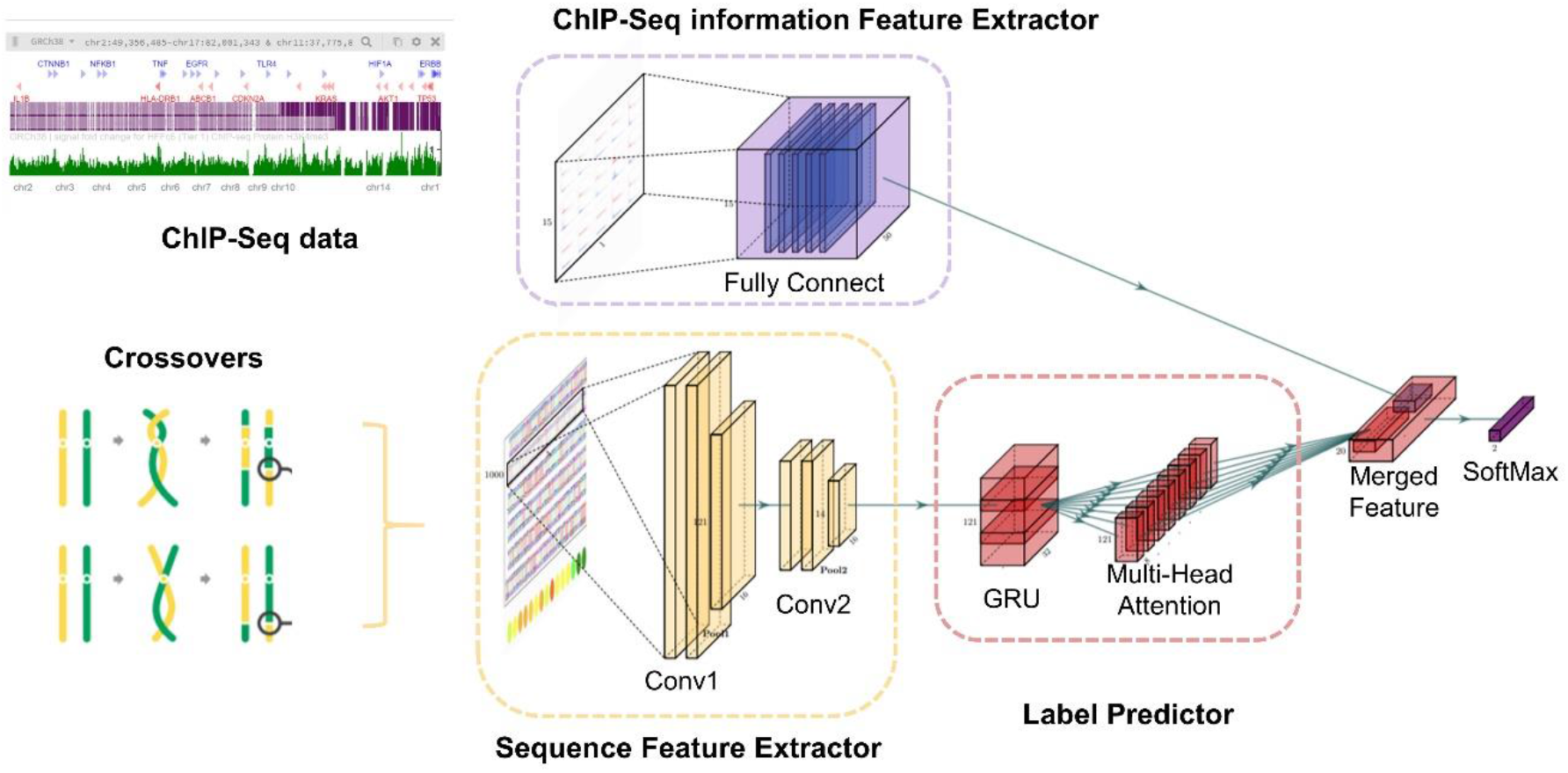
Deep Learning network architecture of RHSNet-chip. The detailed identification model of our proposed RHSNet-chip framework. The input sequence would first go through two 1-D convolutional layers as the sequence feature extractor. Then it will go through a Gated Recurrent Unit (GRU) for capturing long-range information, and a multi-head attention layer for detecting interactions within the sequence. In parallel, the ChIP-seq information would go through a fully connected network. Finally, the sequence feature and the ChIP-seq feature would be merged and give out the final prediction via the SoftMax activation.

**Fig. S3.**
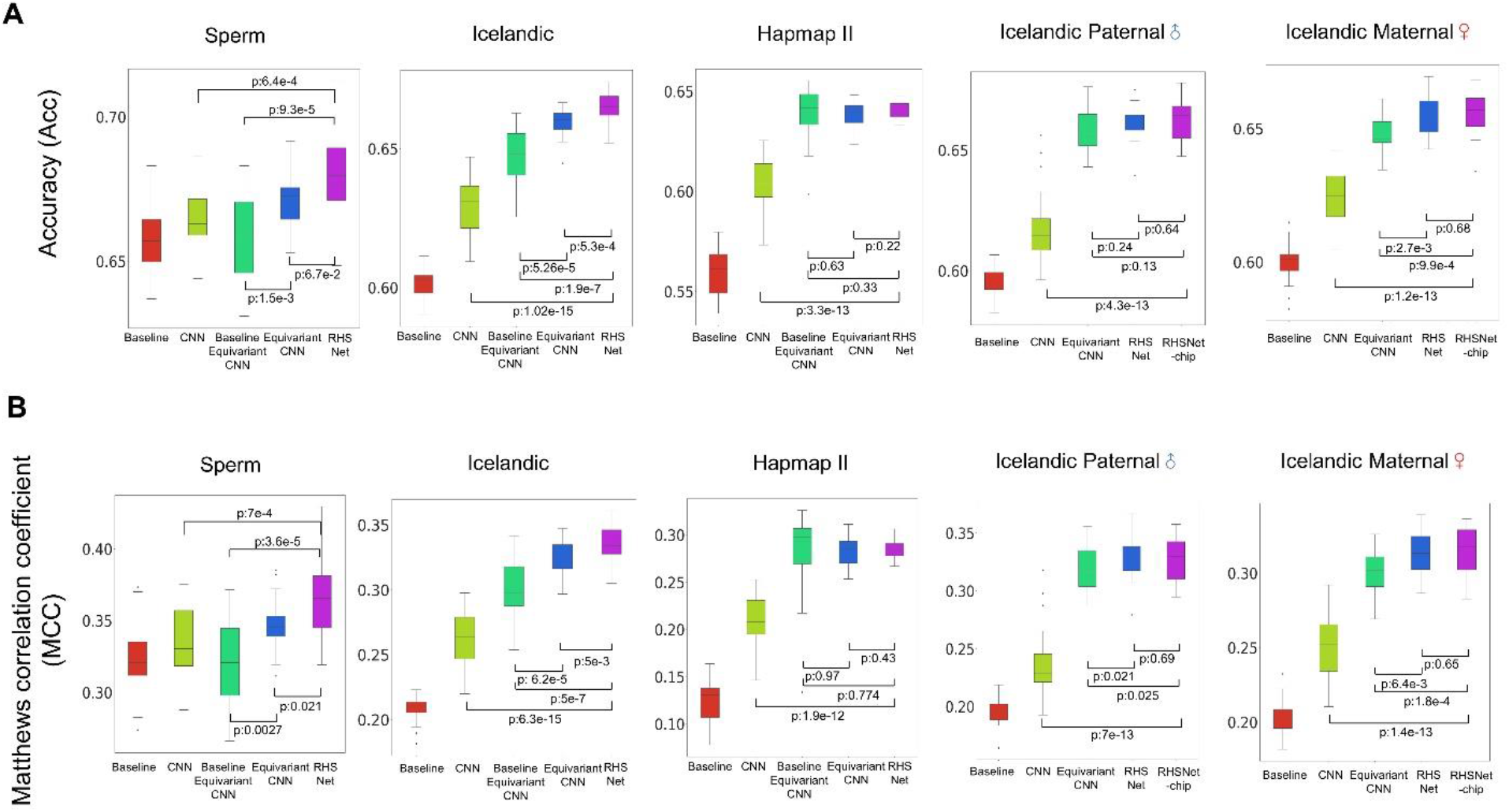
Detailed data statistical performance of RHSNet across different studies and sexes. **(A)** Boxplot of accuracy (Acc) distribution that distinguishes RHSNet from Baseline CNN and Equivariant CNN(*52*) in multiple trials of 5-fold cross-validation experiments. **(B)** Boxplot of Matthews correlation coefficient (MCC) distribution that distinguishes RHSNet from Baseline CNN and Equivariant CNN in multiple trials of 5-fold cross-validation experiments.

**Fig. S4.**
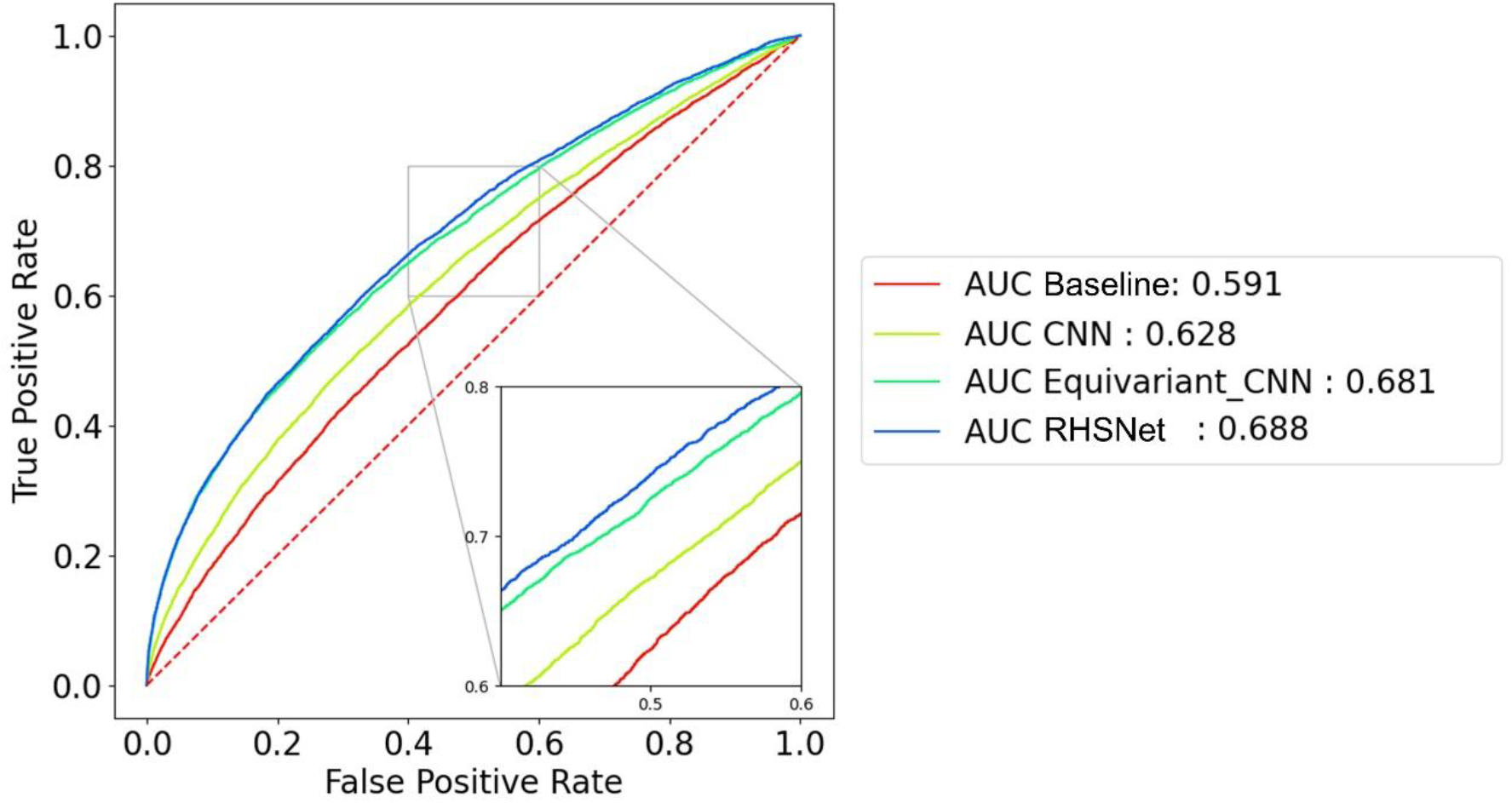
Imbalanced test result on Icelandic dataset. In the Icelandic 2019 dataset, we show the ROC curve and the AUROC score of 4 prediction methods during imbalanced testing. RHSNet achieves the best prediction performance with an AUROC score of 0.688.

**Fig. S5.**
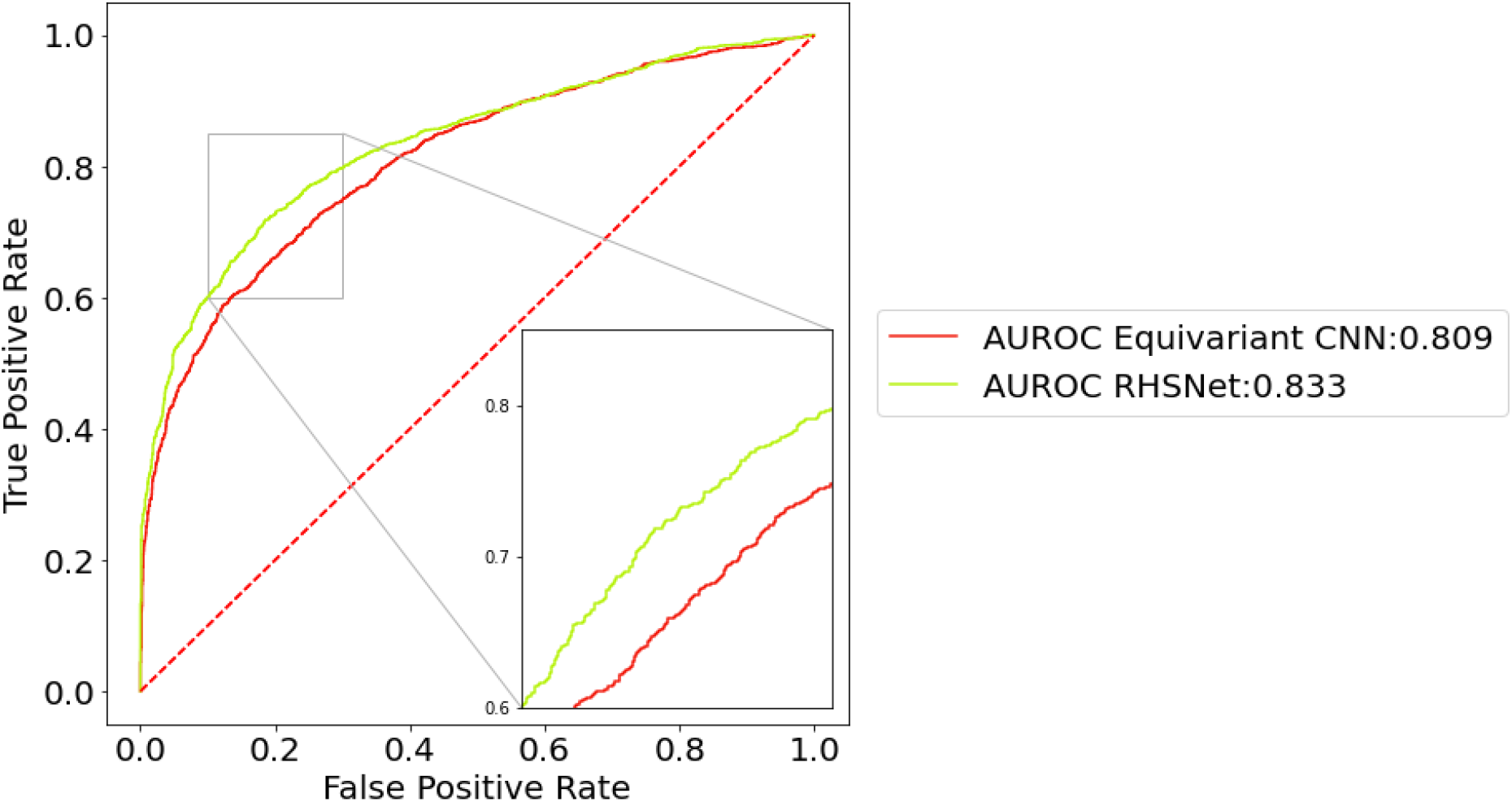
Head-to-head comparison against Equivariant CNN(*52*) on the Yeast dataset. In the Icelandic 2019 dataset, we show the ROC curve and the AUROC score of 4 prediction methods during imbalanced testing. RHSNet achieves the best prediction performance with an AUROC score of 0.688.

**Fig. S6.**
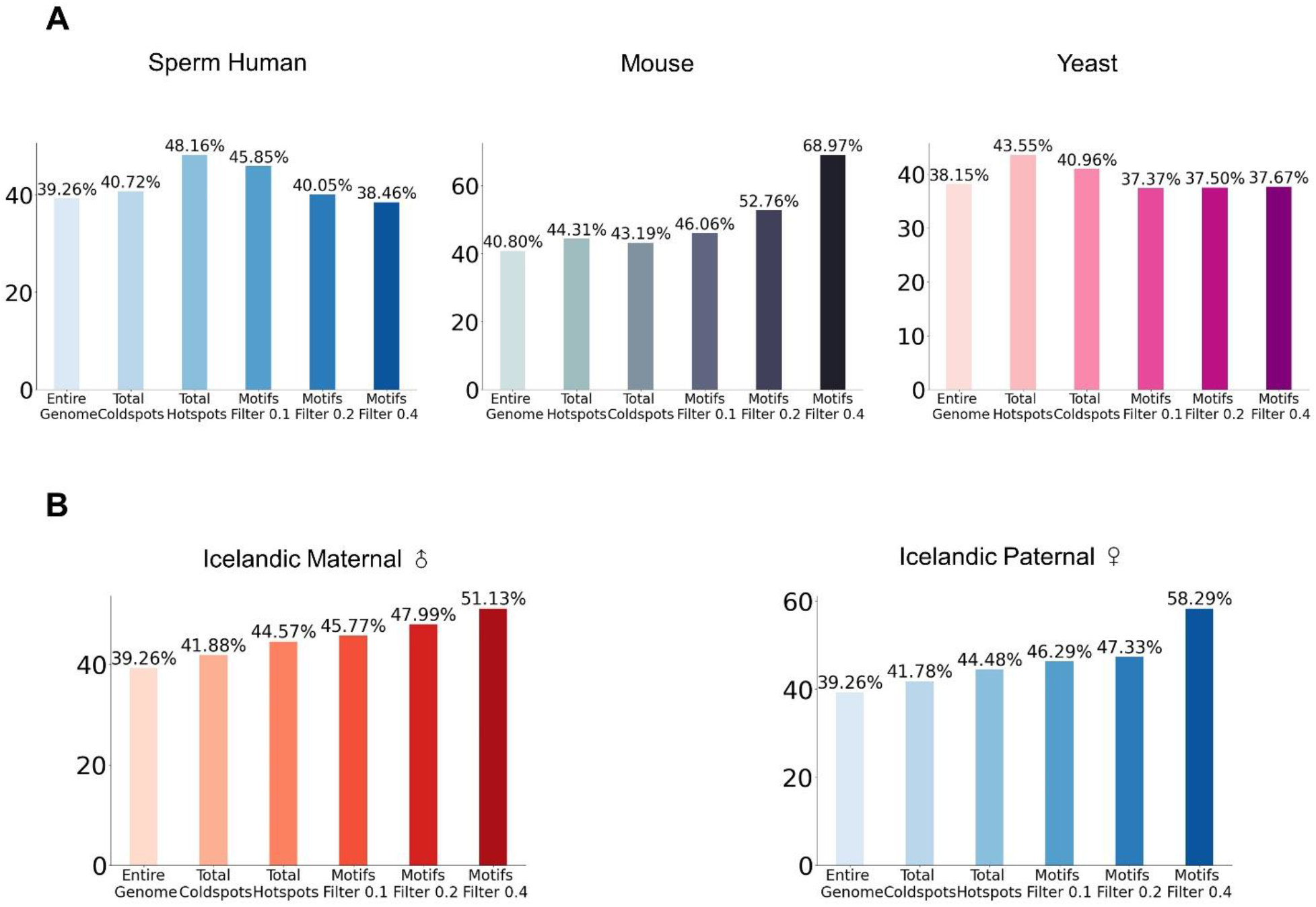
Supplementary GC content results across different species and sexes. **(A)** Statistical comparison of GC content across different species. **(B)** Statistical comparison of GC content across different sexes. The result is calculated from Icelandic(*53*) Human genetic maps.

**Fig. S7.**
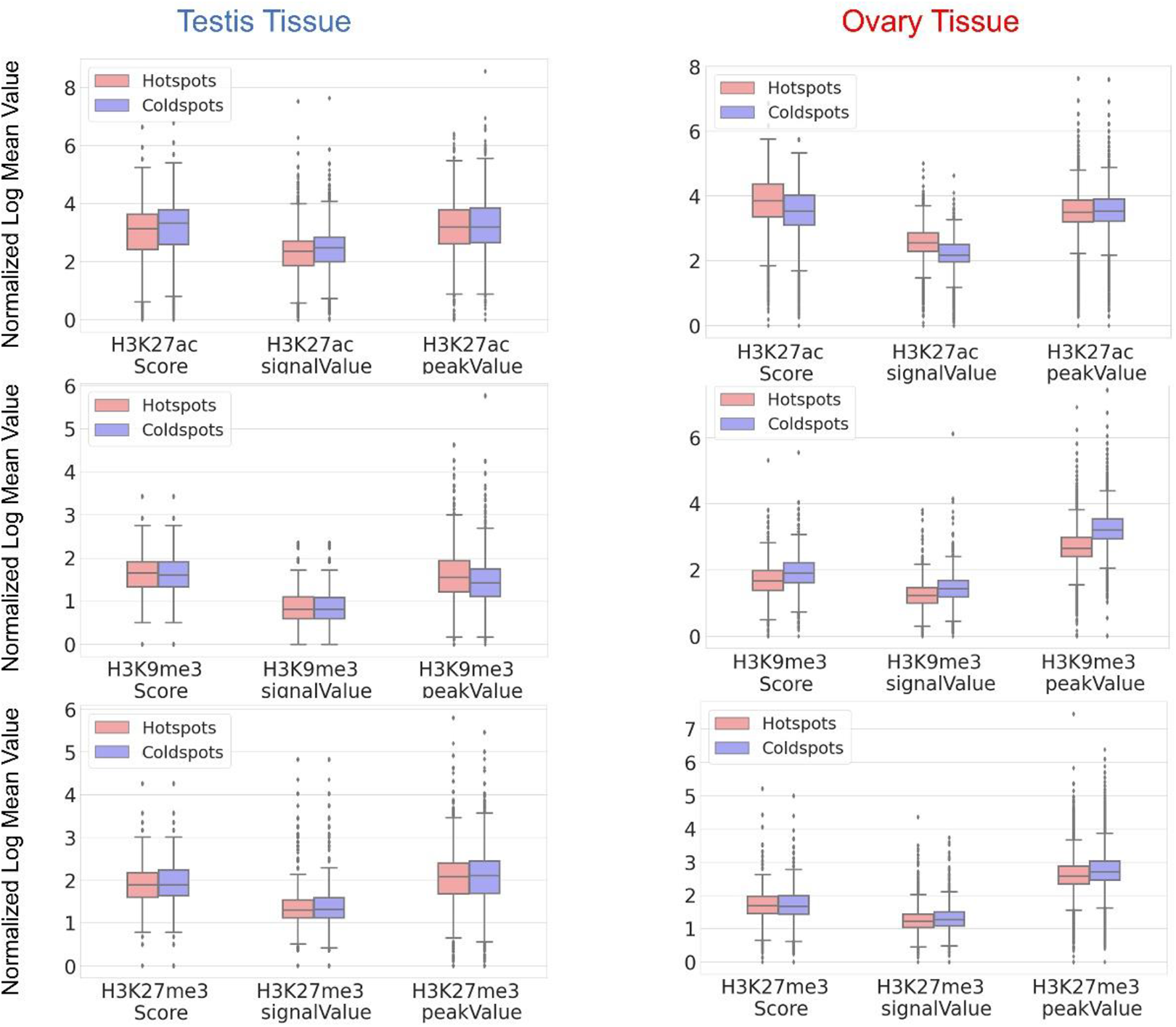
Histone modification feature distribution comparison on ovary tissue between hotspots and coldspots on maternal recombination map. The normalized log mean value comparing the different distributions between hotspots and coldspots from the ChIP-seq features (signal Value, peak Value, Score) extracted from the ovary and testis tissues of Homo sapiens female and male adults. Here, we show the comparison over three different histone modifications: H3K27ac, H3K27me3, and H3K9me3.

**Fig. S8.**
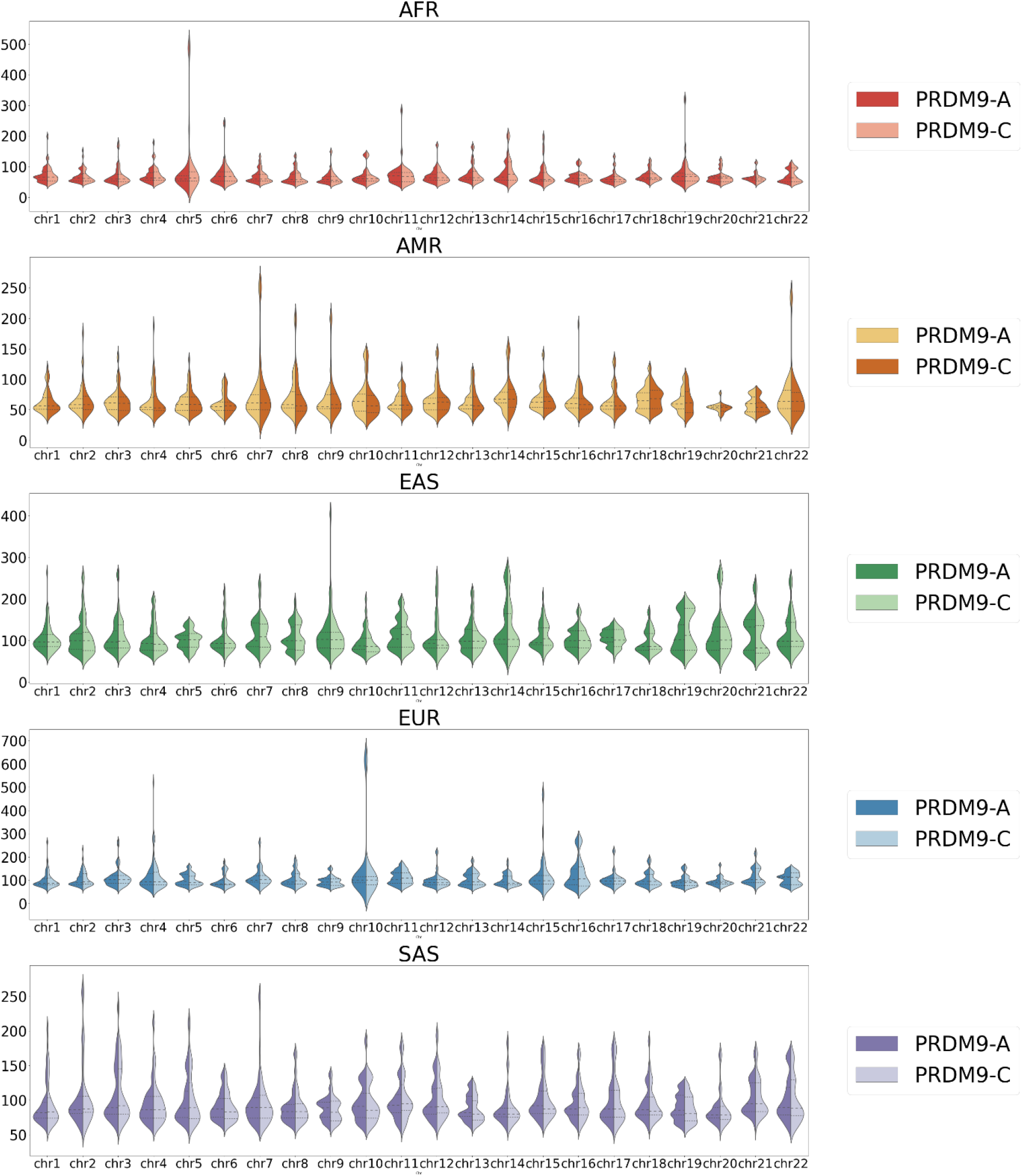
Recombination rate distribution over 22 regular chromosomes across populations. Across five different populations from the 1000 Genome Project(*18*), we draw the recombination rate distribution of PRDM9-A/C alleles over 22 regular chromosomes on the ground truth genetic maps.

**Fig. S9.**
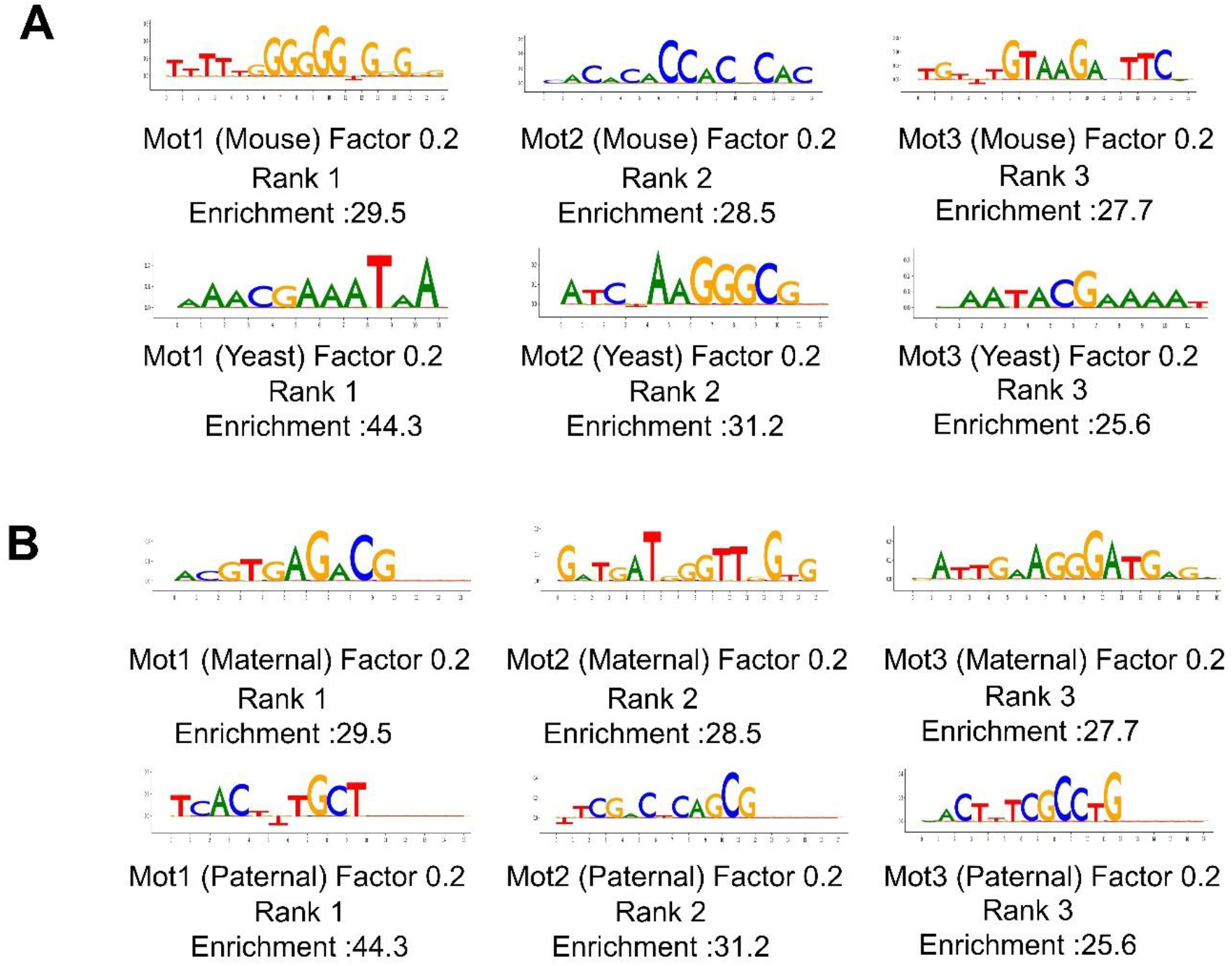
The most important motifs detected in different species and sexes. **(A)**The top 3 detected motifs from the Mouse(*11*) and Yeast(*6*) datasets. **(B)**The top 3 detected motifs from the paternal and maternal genetic maps, respectively.

**Fig. S10.**
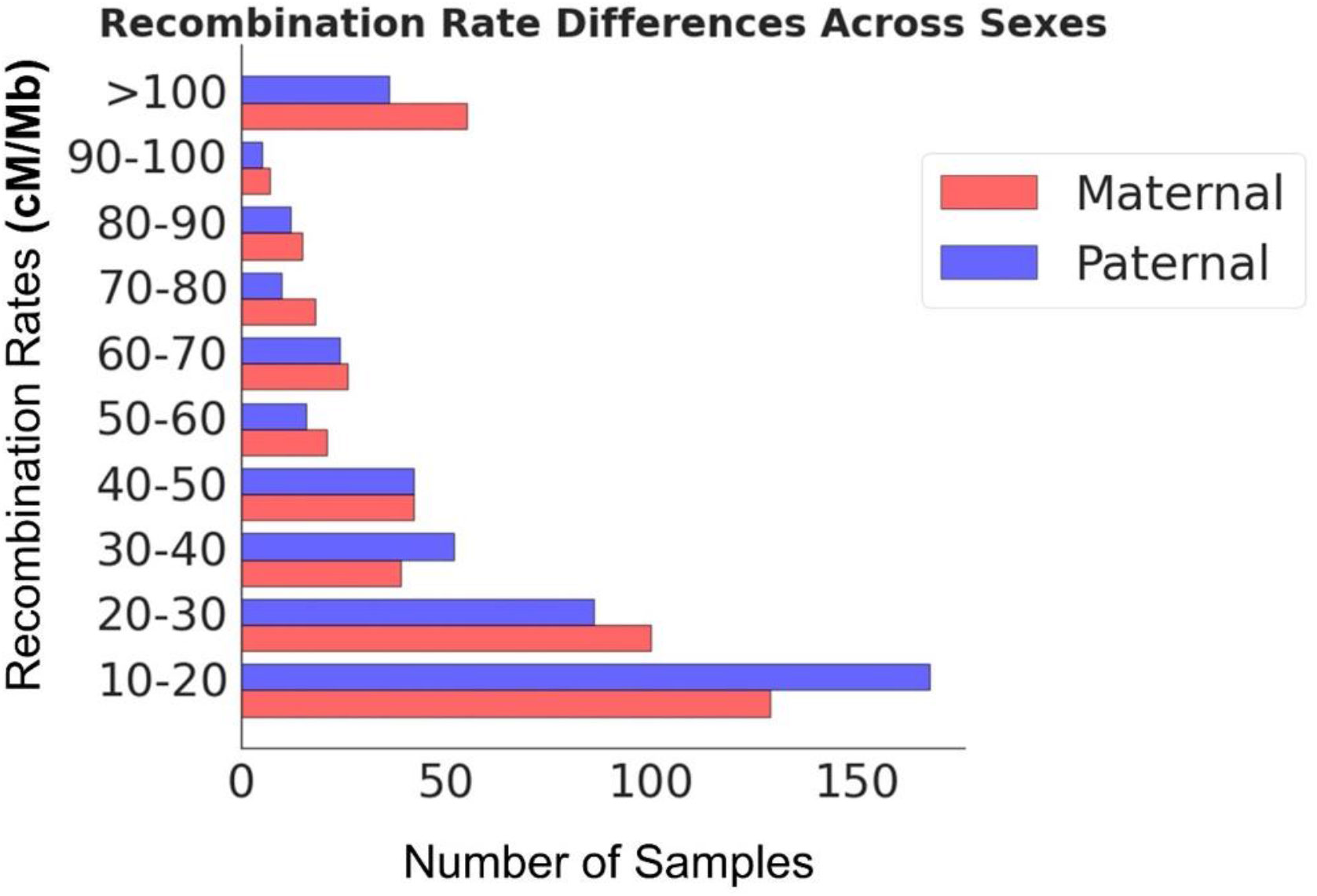
Recombination rate comparison between paternal and maternal motifs. Statistical comparison of the recombination rate between the detected paternal and maternal motifs.

**Fig. S11.**
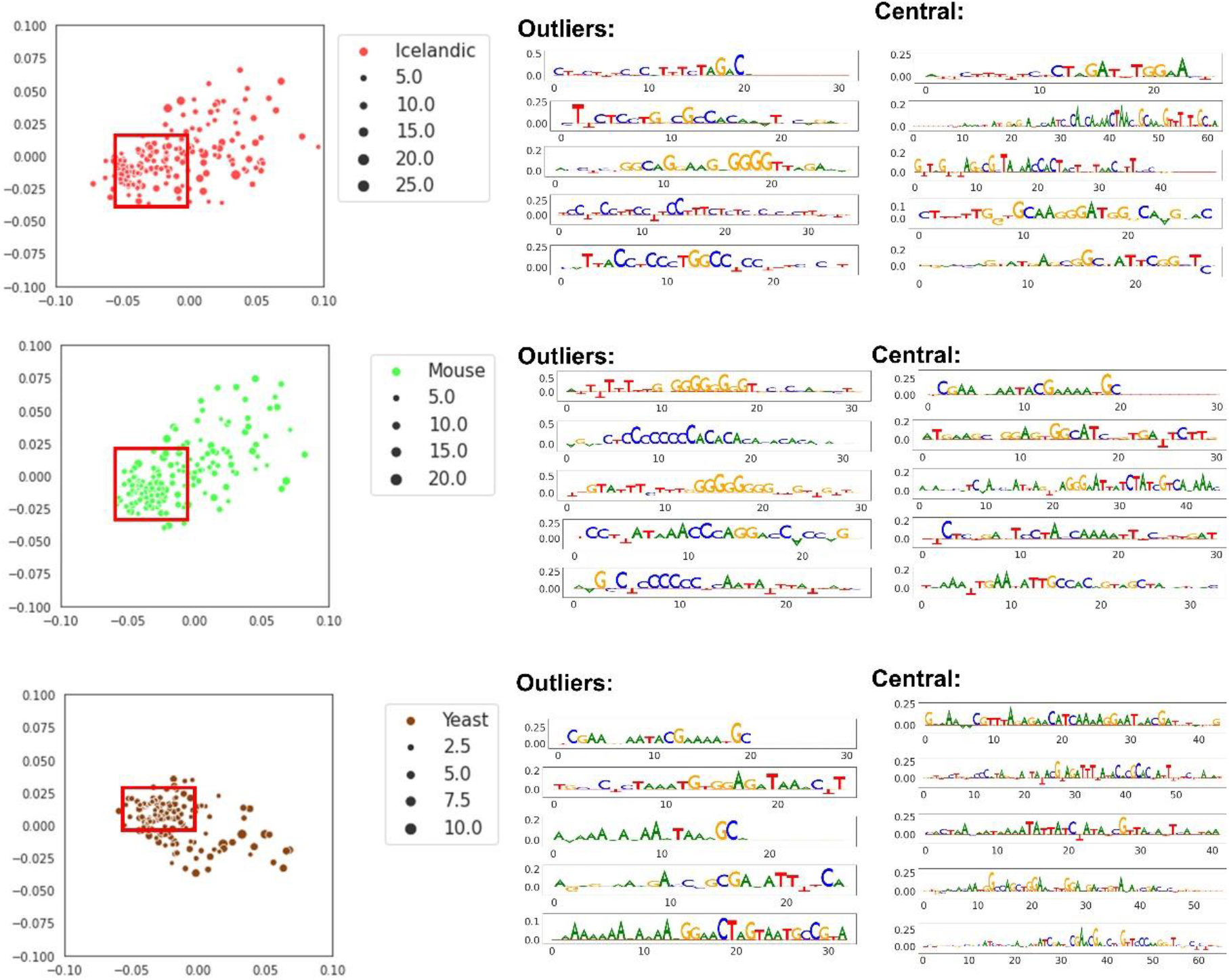
Detailed motif embedding results across species. Embedding vectors in 2D space across different species. The central region is bounded by a red bounding box, and the outliers are defined by comparing the local density of a sample to the local densities of its 50 neighbors. The top 5 ranked motifs are visualized for both central and outlier motifs.

**Fig. S12.**
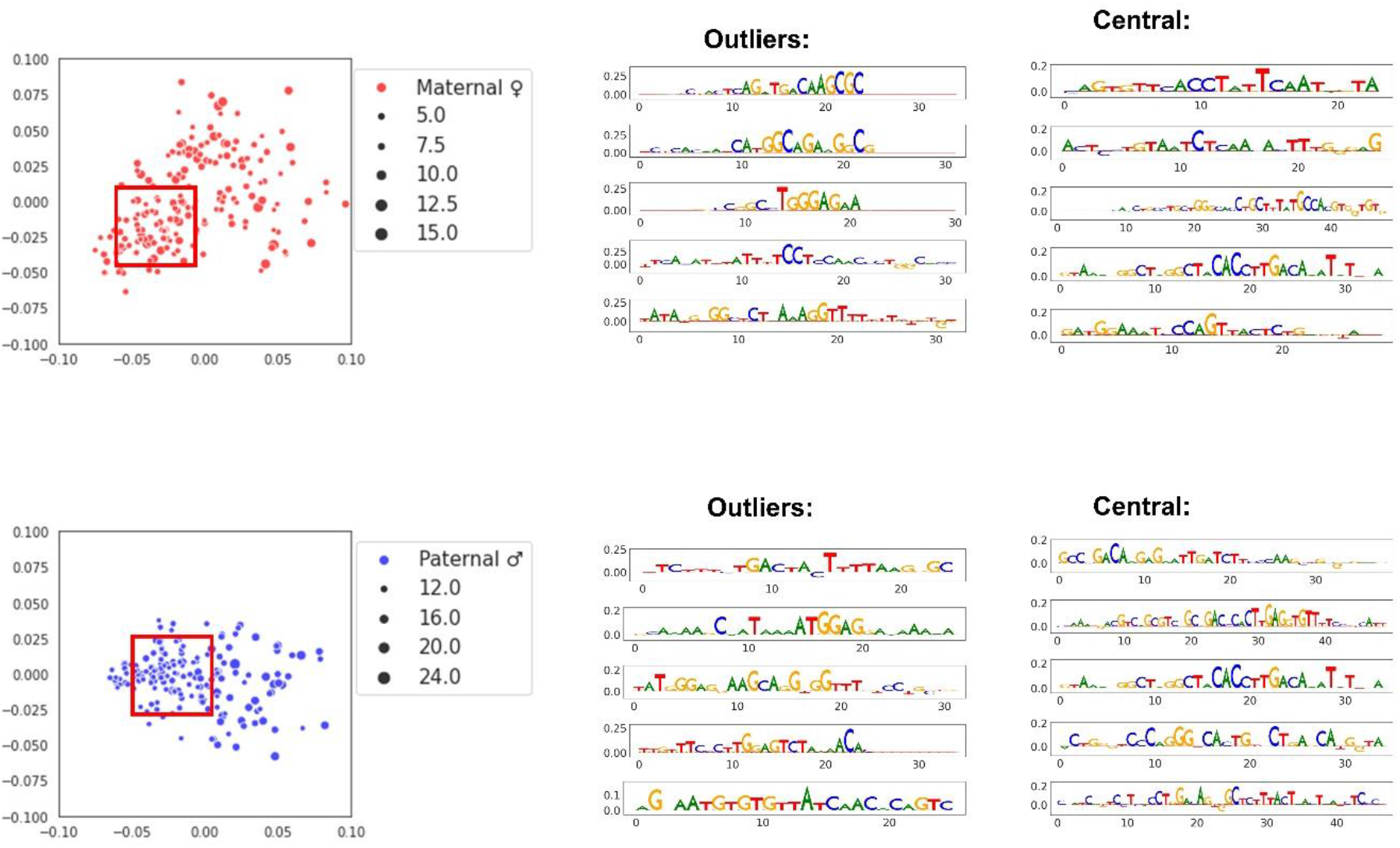
Detailed motif embedding results across sexes. Embedding vectors in 2D space across different sexes. The central region is bounded by a red bounding box, and the outliers are defined by comparing the local density of a sample to the local densities of its 50 neighbors. The top 5 ranked motifs are also visualized for both central and outlier motifs.

**Fig. S13.**
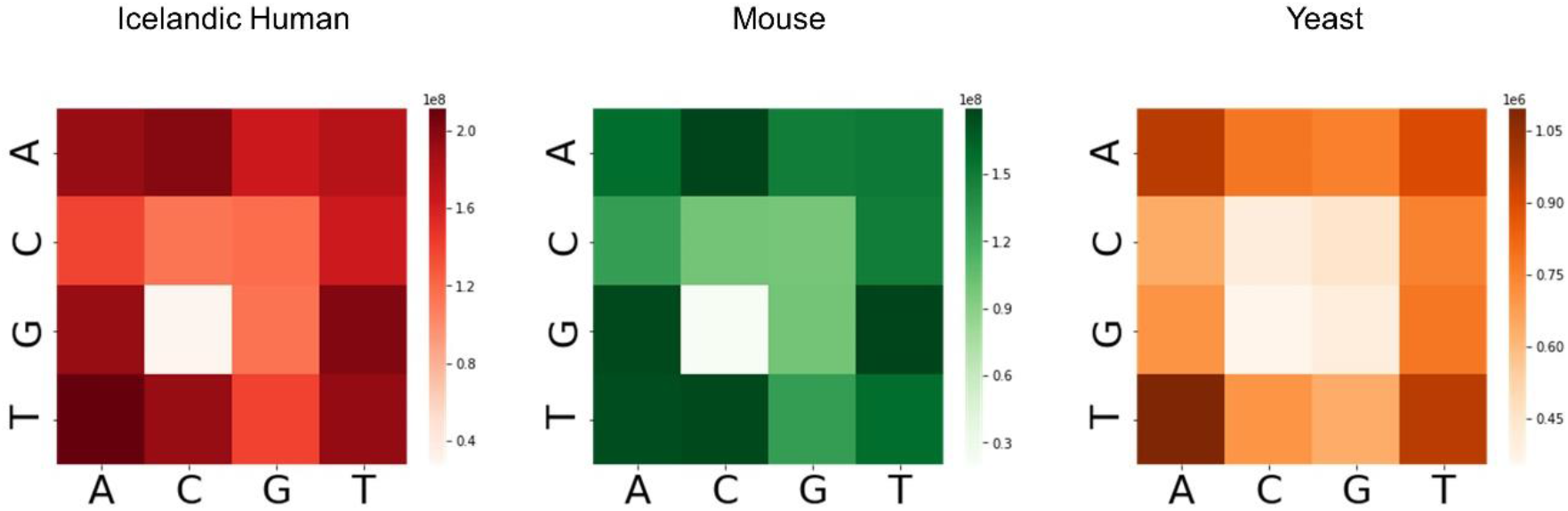
2-mer heatmap visualization of the entire genome. Heatmaps of 2-mer distributions over the entire genome. Each grid represents the 2-mer appearance frequency (AA, AC, AG, AT, CA, CC, CG, CT, GA, GC, GG, GT, TA, TC, TG, TT) across GRCh38 human reference genome, GRCm38/mm10 reference genome, and S288C Yeast reference genome.

**Fig. S14.**
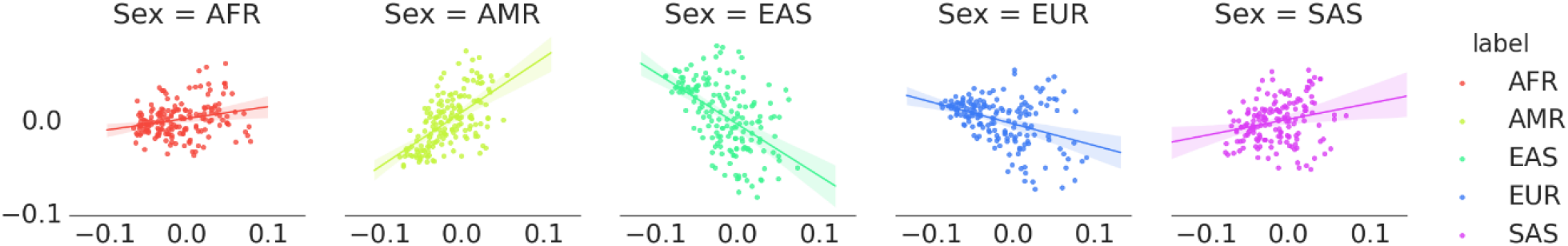
Linearly regressed embedding results across populations. The embedding results with regression models for five different populations.

**Fig. S15.**
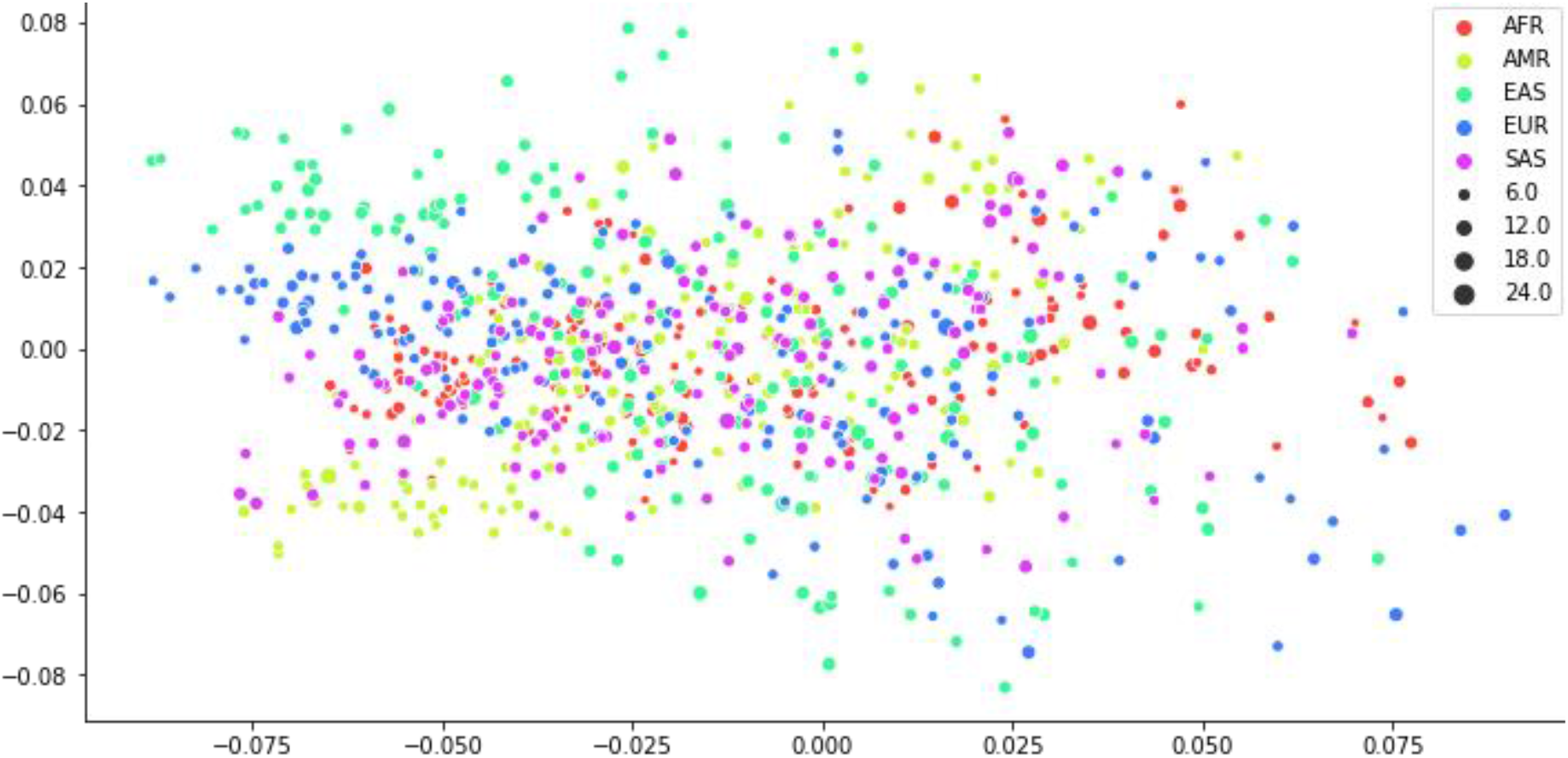
Embedding results across populations. The overall embedding results within the same 2D space for five different populations.

**Table S1.** Statistics of recombination hotspot dataset across different studies. Statistical comparison of our hotspots/coldspots construction across different studies on the human genome, with an imbalanced testing dataset on Icelandic 2019 dataset. The average recombination rate of each study is attached under each row.

|  | 5-fold Cross-validation |  | Imbalanced Testing |  |
| --- | --- | --- | --- | --- |
|  | Hotspots | Coldspots | Hotspots | Coldspots |
| Icelandic<br>2019(53) | 20,000 | 20,000 | 5,467 | 95,733 |
|  | 51.07cM/Mb | 1.78e10-10<br>cM/Mb | 43.18cM/Mb | 0.07 cM/Mb |
| HapMap II<br>2008(54) | 17,552<br>10.5cM/Mb | 17,547<br>0.5cM/Mb | —— | —— |
| Sperm 2020(13) | 5,000<br>19.96cM/Mb | 5,000<br>1e10-20 cM/Mb | —— | —— |

**Table S2.** Statistics of recombination hotspot dataset across different species and sexes. Statistical comparison of dataset construction across different sexes and different species. The average recombination rate of each study is attached under each row.

|  | Hotspots | Coldspots |
| --- | --- | --- |
| Icelandic Paternal ♂ | 15,000<br>44.13cM/Mb | 15,000<br>1.2e10-14cM/Mb |
| Icelandic Maternal ♀ | 20,000<br>48.28cM/Mb | 20,000<br>1.8e10-11cM/Mb |
| Mouse(*11*) | 9,620 | 9,620 |
| Yeast | 468 | 468 |

**Table S3.** Statistics of recombination hotspot dataset across different populations. Statistical comparison of the 1000 Genome(*18*) dataset construction across five populations with corresponding recombination rate over each population.

| Population Code | Population | Hotspots | Coldspots |
| --- | --- | --- | --- |
| AFR | African | 50,049<br>27.45cM/Mb | 50,035<br>0.0378cM/Mb |
| AMR | Admixed American | 18,160<br>27.32cM/Mb | 18,152<br>0.0390cM/Mb |
| EAS | East Asian | 27,030<br>38.44cM/Mb | 27,020<br>0.094cM/Mb |
| EUR | European | 31,283<br>35.50cM/Mb | 81,273<br>0.0380cM/Mb |
| SAS | South Asian | 32,593<br>33.91cM/Mb | 32,583<br>0.0383cM/Mb |

**Table S4.**
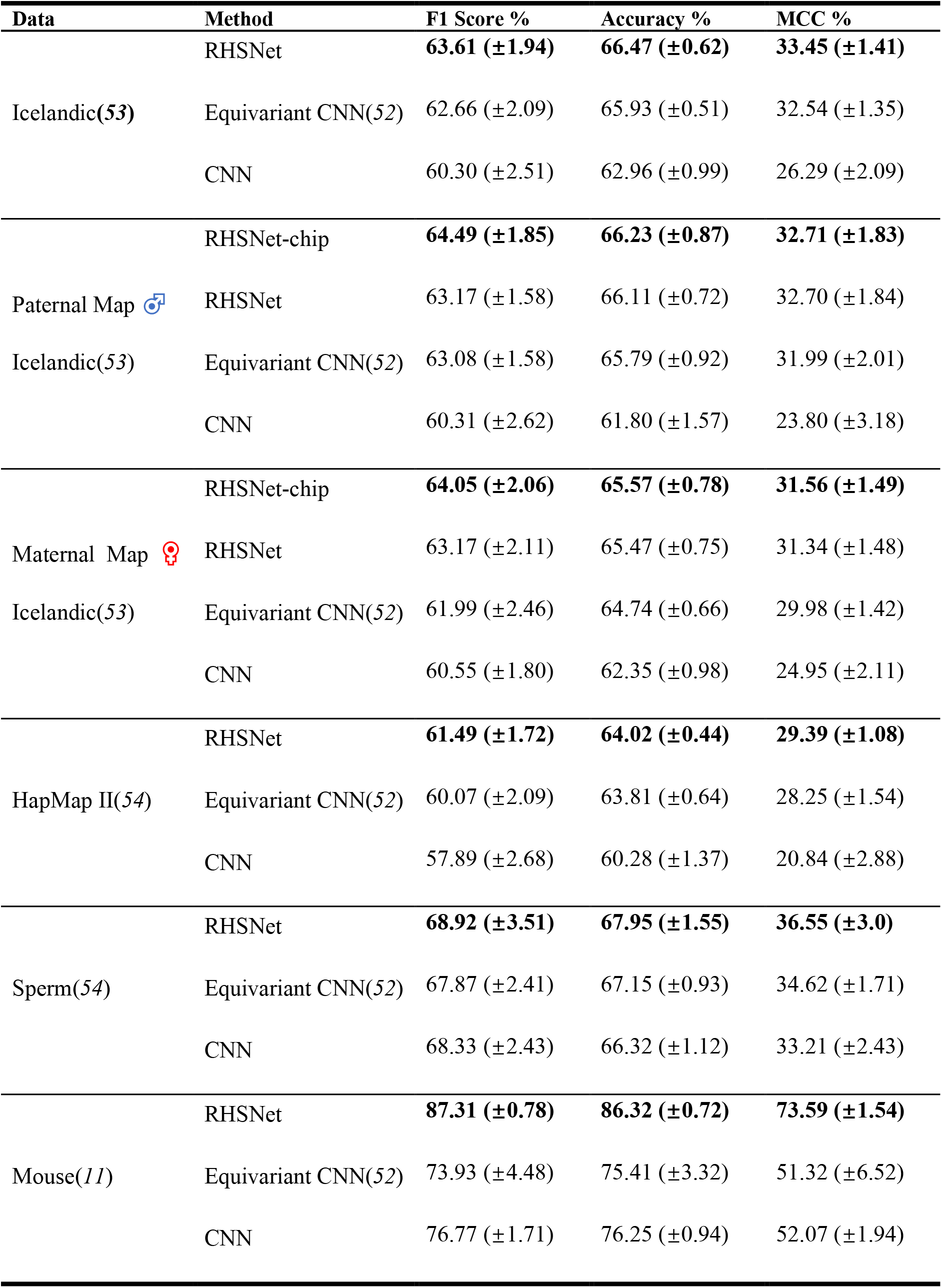
Detailed overall classification performance statistics on different datasets. Detailed classification performance of the proposed RHSNet-chip and RHSNet, compared with the baseline CNN model and the existing Equivariant CNN(*52*). RHSNet shows outstanding prediction performance on multiple benchmark datasets: Icelandic(*53*), HapMap II(*54*), Sperm(*13*), and Mouse(*11*), which are across different studies, sexes, and species.

**Table S5.** Comparison of RHSNet against shallow learning methods. Statistical results on the HapMap II(*54*) dataset, comparing the proposed RHSNet with another bassline method: PseAAC(*55*) + Support Vector Machine (SVM(*56*)) classifier. Multiple experiments with different sets of parameter lambda for pseudo-feature extraction are conducted. The deep learning-based methods show a significant edge over the SVM-based classifier.

| Method | lambda | weight | Overall Accuracy% |
| --- | --- | --- | --- |
| RHSNet | -- | -- | 64.02 |
| Equivariant CNN | -- | -- | 63.55 |
| Baseline | -- | -- | 60.28 |
| PseAAC + SVM | 3 | 0.05 | 54.24 |
| PseAAC + SVM | 5 | 0.05 | 53.96 |
| PseAAC + SVM | 10 | 0.05 | 54.61 |
| PseAAC + SVM | 20 | 0.05 | 54.92 |

**Table S6.**
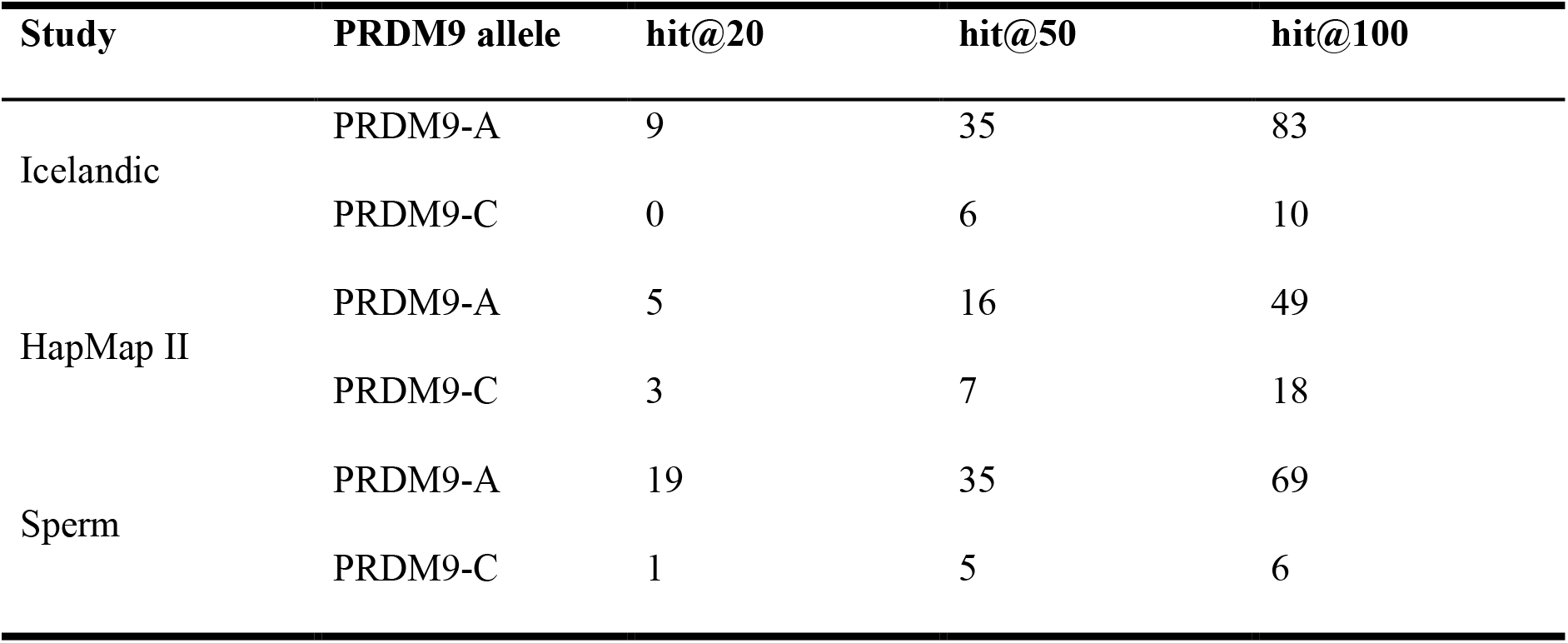
PRDM9 alleles identification across different studies. The hit@20/50/100 results of the RHSNet’s identification across different studies.

**Table S7.** PRDM9 alleles identification across different populations. The hit@20/50/100 results of the RHSNet’s identification across different populations.

| Population | PRDM9 allele | hit@20 | hit@50 | hit@100 |
| --- | --- | --- | --- | --- |
| African | PRDM9-A | 20 | 58 | 124 |
|  | PRDM9-C | 0 | 0 | 8 |
| American | PRDM9-A | 19 | 25 | 44 |
|  | PRDM9-C | 0 | 4 | 12 |
| East Asian | PRDM9-A | 0 | 2 | 7 |
|  | PRDM9-C | 0 | 2 | 3 |
| European | PRDM9-A | 14 | 24 | 48 |
|  | PRDM9-C | 0 | 3 | 4 |
| South Asian | PRDM9-A | 30 | 71 | 90 |
|  | PRDM9-C | 0 | 2 | 2 |

